# Fishash: A contingency table approach to Perturb-seq guide assignment

**DOI:** 10.64898/2026.01.22.701179

**Authors:** Jack Kamm, Jake Yeung, William F. Forrest

## Abstract

**Background:** Single-cell pooled CRISPR screens (Perturb-seq) are a powerful tool in functional genomics. A key preprocessing step is to determine which cells received which perturbations based on possibly noisy sequencing counts of the guide RNA library. Many existing approaches to this problem require fitting probabilistic models which may be computationally expensive on large screens with 10s of thousands of cells and guides.

**Results:** We propose to view the guide count matrix as a contingency table and use Fisher’s Exact Test to test for associations between cell and guide barcodes. This approach is fast, normalizes for both cell and guide-specific size factors, and provides a p-value for each cell-guide pair. Our method further uses a multiple testing correction approach that accounts for the correlation structure between the tests, and a correction for Simpson’s paradox that arises due to hidden confounding. Additionally, to facilitate the development and benchmarking of guide assignment methods, we propose a framework for simulating guide counts with a realistic model of sequencing noise.

**Conclusions:** We find that our method compares favorably to existing methods in both accuracy and runtime on simulated and real datasets. We provide our method in an easy to use R package, fishash, available at https://github.com/jackkamm/fishash. Additionally, the code to reproduce the results of this manuscript is available at https://github.com/jackkamm/fishash_analysis.

## 1 Background

Single-cell pooled CRISPR screens (Perturb-seq) have emerged as a powerful tool in functional genomics to understand the consequences of genetic perturbations on downstream phenotypes (Dixit et al., 2016; Datlinger et al., 2017; Replogle et al., 2022). A key preprocessing step to analyze this type of data is to determine which cells received which CRISPR perturbations, based on sequencing read counts of the guide RNA library. However, different sources of noise can cause contaminating reads, for example due to extracellular debris from lysed cells or barcode swapping due to PCR chimeras (Dixit, 2016; Fleming et al., 2023). Therefore, it is critical to determine which observed cell-guide pairs had reads due purely to noise. Many existing methods use probabilistic mixture models to infer which reads are from signal or noise (Liu et al., 2025; Braunger and Velten, 2024; Barry et al., 2024), but fitting these models can be computationally expensive, especially on genome-wide Perturb-seq screens with tens of thousands of guides and millions of cells (Replogle et al., 2022).

We propose to assign guides using a classical approach from the analysis of contingency tables, by treating the UMI count matrix as a contingency table and examining its association factors (Good, 1956), which are the odds ratios for all of its 2 × 2 sub-tables. More specifically, for each cell barcode and guide barcode, we test whether the co-occurrence of the 2 barcodes has an odds ratio greater than 1 using a one-sided Fisher’s exact test (Fisher, 1922). This approach is fast, and performs surprisingly well, outperforming more sophisticated and computationally intensive mixture models on both real and simulated data. While writing this manuscript, we learned that a similar approach had also been proposed by Teyssier (2024), implemented in the Python package geomux.

However, using Fisher’s exact test on its own has two drawbacks: first, Simpson’s paradox can cause false associations between cell and guide barcodes due to hidden confounding (Simpson, 1951); and second, the test statistics from Fisher’s exact test are negatively associated, which invalidates FDR control from the Benjamini-Hochberg method (Benjamini and Hochberg, 1995; Benjamini and Yekutieli, 2001). We address the first issue by characterizing the conditions under which Simpson’s paradox will cause false associations, and by proposing a correction for Simpson’s paradox that involves iteratively estimating signal and noise components and re-computing Fisher’s test. We address the second issue by providing options for multiple testing procedures that account for dependency; by default, fishash uses the procedure of Guo and Sarkar (2020), which controls FWER and FDR assuming independence across cells but dependence for guides within a cell, which is a reasonable approximation for most Perturbseq datasets, and provides a middle ground between the Benjamini-Hochberg and Benjamini-Yekutieli procedures.

We additionally propose a framework for simulating guide counts using a realistic model of sequencing noise (Fleming et al., 2023). We anticipate this simulation framework will be useful for future development and bench-marking of guide assignment methods.

Using simulated and real datasets (Liu et al., 2025; Replogle et al., 2022), we demonstrate that our method is powerful, computationally efficient, and does a better job of controlling type I error at specified levels compared to other methods in many scenarios. We provide our method in a fast and easy to use R package, fishash.

## 2 Methods

### 2.1 One-sided Fisher test

For UMI *u* = 1, …, *n*_umi_, let *C*_*u*_ ∈ C be its cell barcode and *G*_*u*_ ∈ G be its guide barcode. Let

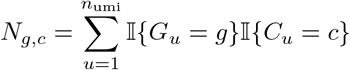

be the number of UMIs with barcode pair (*g, c*). Define *N*_*g*,:_ and *N*_:,*c*_ be the total guide and cell UMI counts respectively,

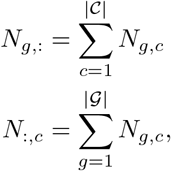

and let *N*_−*g,c*_, *N*_*g*,−*c*_, *N*_−*g*,−*c*_ correspond to the total counts without barcodes *g* and/or *c*:

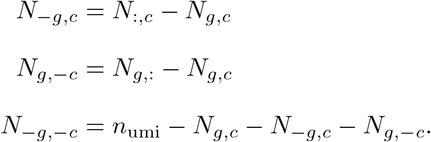

Let *π*_*g,c*_ denote the true frequency of barcode pair (*g, c*) if we sequenced the reads at infinite sequencing depth, so that (*N*_*g,c*_) ~ Multinomial(*n*_umi_, (*π*_*g,c*_)) (see also de Finetti’s Theorem). Analogous to above, we define the terms

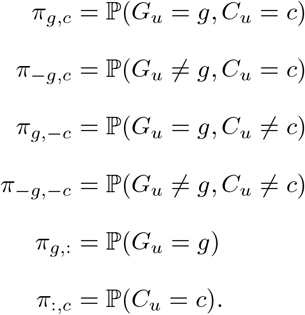

For the 2 × 2 contingency table

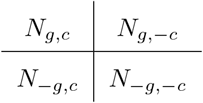

let *R*_*g,c*_ be the latent odds-ratio and 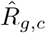 be the empirical odds-ratio,

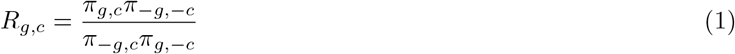

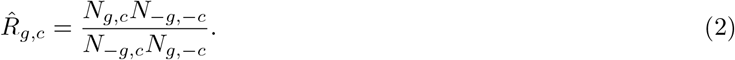

For each (*g, c*), we test the null hypothesis 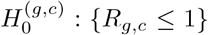 against the alternative 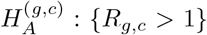, and to that end define *P*_*g,c*_ as the p-value under a one-sided Fisher’s exact test

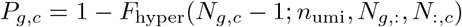

where *F*_hyper_(·; *N, K, n*) is the CDF of a hypergeometric distribution where *N* is population size, *K* is number of success states in population, and *n* is number of draws.

### 2.2 Correction for Simpson’s paradox

In the context of 2 × 2 contingency tables, Simpson’s paradox (Simpson, 1951) occurs when there is a confounding variable such that the log-odds-ratio flips sign when we condition on i t. In the case of g RNA UMIs, each UMI can be considered as coming from “signal” or “noise”, which is a hidden latent variable. More specifically, a UMI with cell barcode *c* that truly originated from cell *c* would be a signal UMI, whereas a UMI with cell barcode *c* that didn’t originate from *c* would be a noise UMI; such noise UMIs can be generated by a couple different processes, dicussed in Section 2.4. Failing to condition on whether a UMI is from signal or noise can cause a positive association between cell barcode *c* and guide barcode *g*, i.e. log(*R*_*g,c*_) *>* 0, even when cell *c* does not contain *g*; we show an illustration in Figure 2.

**Figure 1:**
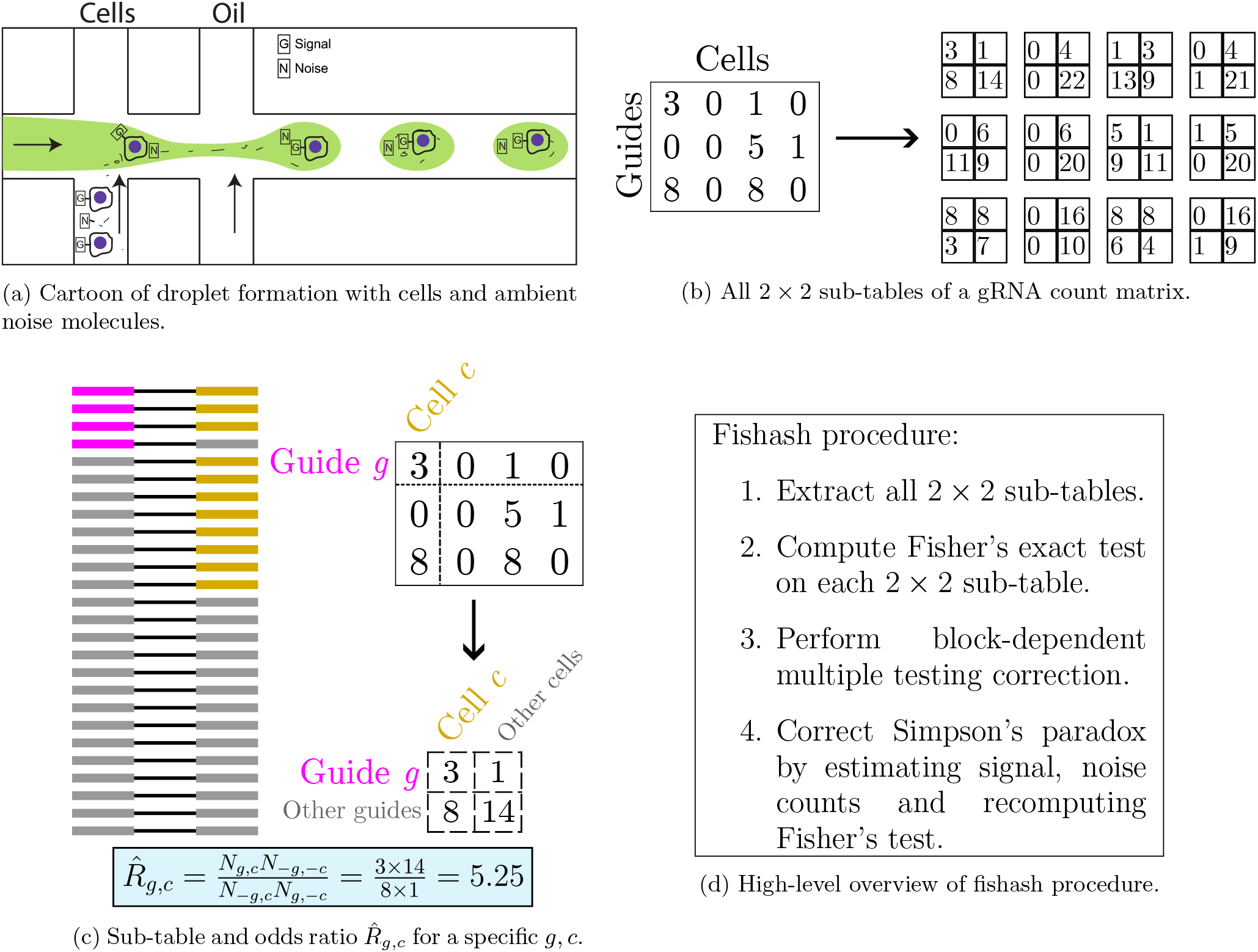
Overview and illustration of the fishash method. The method assigns guides to cells by testing the association between each cell barcode and guide barcode. The method also includes a multiple testing correction that is aware of the block-dependent structure of the tests (Section 2.3), and a correction for false associations from Simpson’s paradox (Section 2.2, Figure 2). See Algorithm 1 for a more complete description of the procedure.

**Figure 2:**
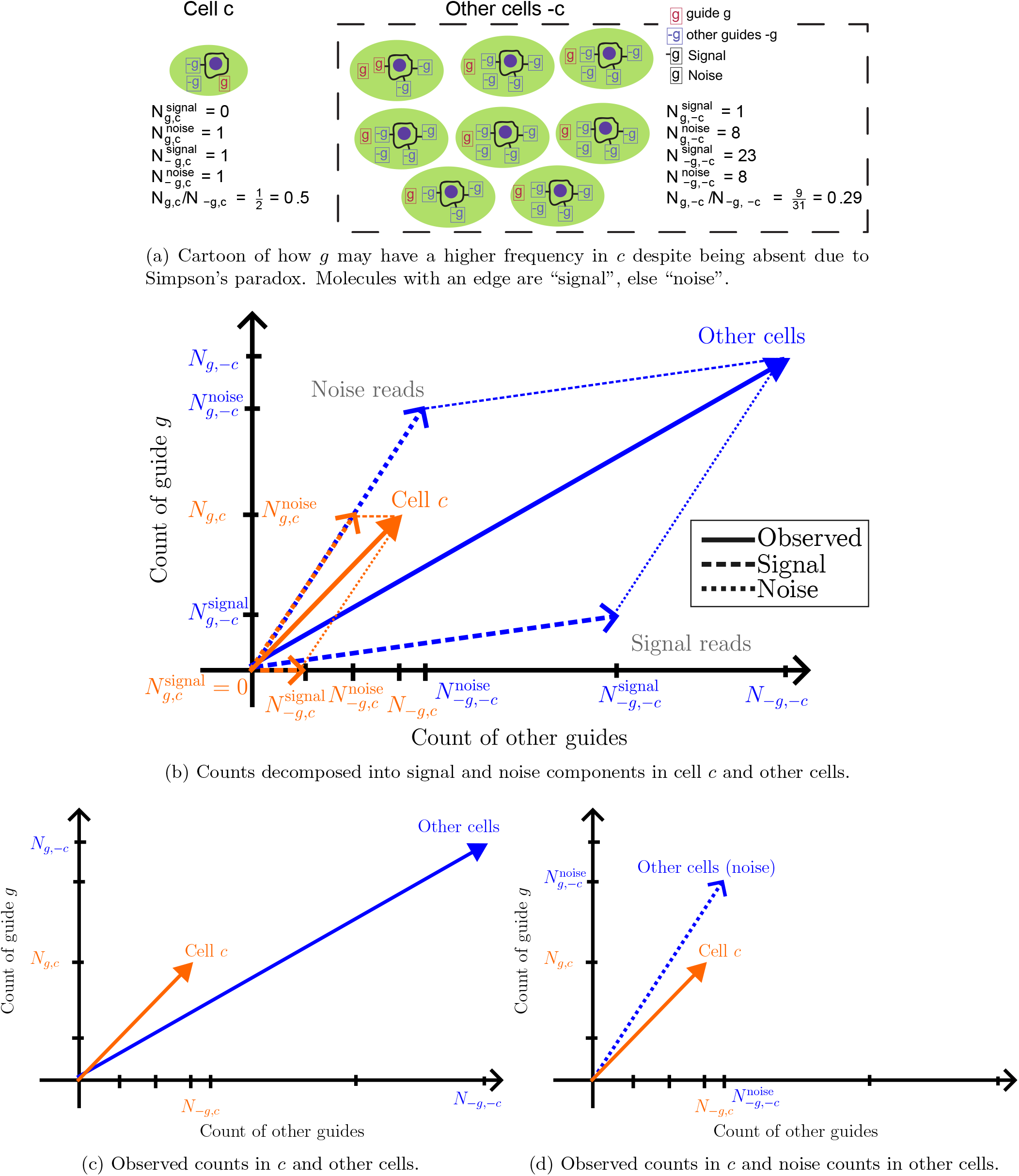
Illustration of how Simpson’s paradox can cause false positives in guides with low signal-to-noise. Figure 2b shows the counts of reads with and without guide barcode *g* and cell barcode *c*, decomposed into signal and noise components. In this example, *g* is not present in *c*, so 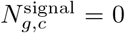. However, due to Simpson’s paradox, barcodes *g* and *c* have odds ratio greater than 1 leading to a false association (Figure 2c, the orange vector has a larger trigonometric tangent than the blue vector). We propose to subtract the signal counts in other cells, which leads to an odds ratio less than 1 and prevents the false association (Figure 2d).

More specifically, let *S*_*u*_ = 1 if the *u*-th UMI comes from the signal, and *S*_*u*_ = 0 if it comes from noise. Let *p*_signal_ = ℙ (*S*_*u*_ = 1) be the fraction of UMIs coming from signal, let *p*_noise_ = 1−*p*_signal_ be the fraction of UMIs coming from the noise, and let 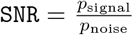 the signal-to-noise ratio. Note that we 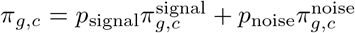, where 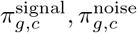 are the probability of (*g, c*) among the signal and noise UMIs respectively:

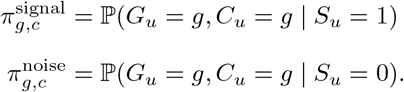

Then the following proposition characterizes when Simpson’s paradox occurs:

#### Proposition 1

*Assume that guide g is not present in cell c, so π*^*signal*^ = 0. *Furthermore, assume that barcodes g and c are independent in the noise reads, so that their joint frequency is the product of their marginal frequencies*,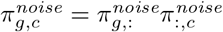. *Then R*_*g,c*_ *>* 1 *if and only if*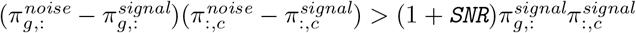.

We prove Proposition 1 in Appendix A. Note that Proposition 1 implies that Simpson’s paradox does not occur whenever 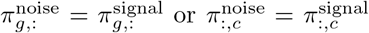– that is, whenever the guide barcode or cell barcode frequencies are equal in the signal and noise UMIs.

Now let 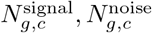 be the counts for (*g, c*) from signal and noise, respectively, so that 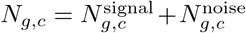. To correct for Simpson’s Paradox, we propose to replace *N*_*g*,−*c*_, *N*_−*g*,−*c*_ in the empirical odds-ratio (2) with the corresponding noise counts 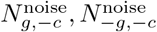 this corresponds to testing an adjusted odds ratio 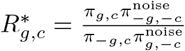. Intuitively, the motivation is to test whether the frequency of guide *g* within cell *c* is above the level of background noise in the other cells (Figure 2d). Since 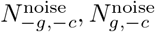 are unobserved, we propose an iterative approach (Algorithm 1), whereby we first perform a standard Fisher test, then generate estimates of the noise counts 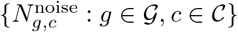, and then recompute Fisher’s test but substituting 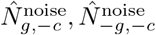 instead of *N*_*g*,−*c*_, *N*_−*g*,−*c*_. This process can be iterated until convergence. We present details of this algorithm, as well as additional mathematical motivation for the adjusted odds ratio 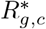, in Appendix B.

#### Algorithm 1

Fishash algorithm.

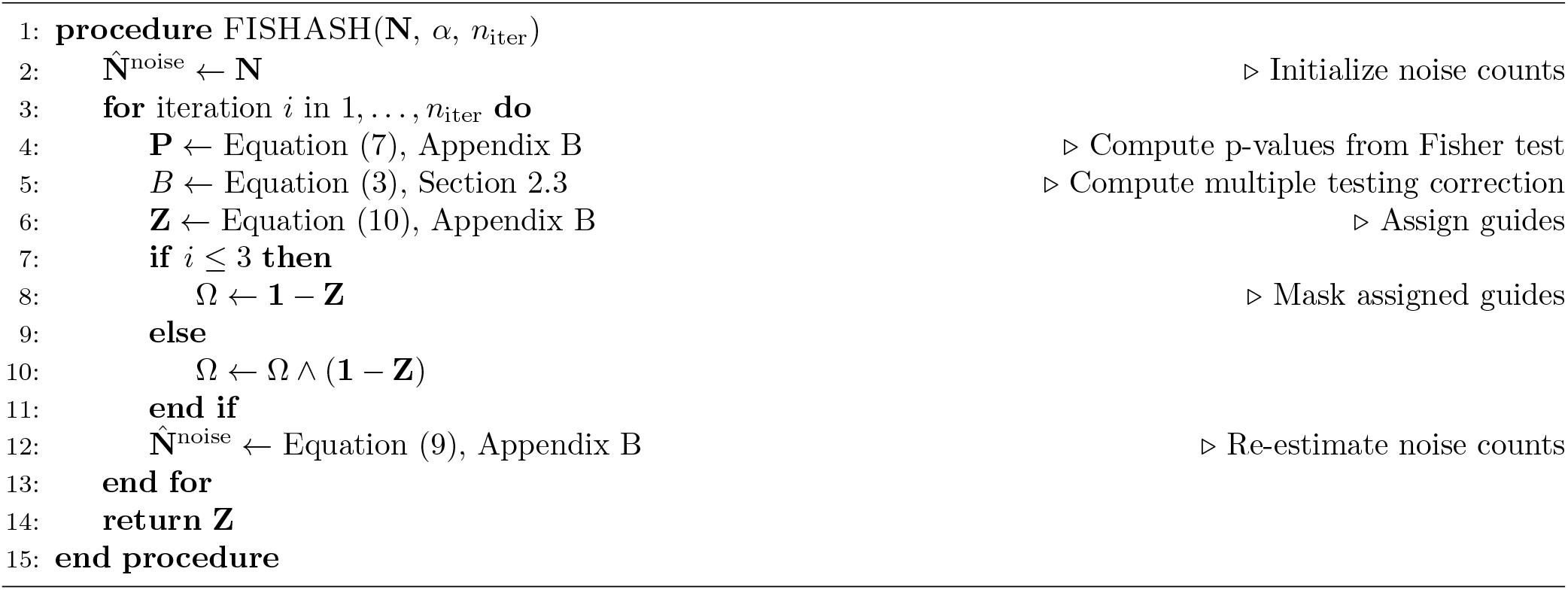

### 2.3 Multiple testing correction

The Benjamini-Hochberg procedure (Benjamini and Hochberg, 1995) is commonly used to correct p-values to control the False Discovery Rate, but requires that the test statistics satisfy Positive Regression Dependency on each one from a Subset, or PRDS (Benjamini and Yekutieli, 2001), a concept related to positive association between variables (Lehmann, 1966; Sarkar, 1969). However, the components of a multivariate hypergeometric distribution are negatively associated (Joag-Dev and Proschan, 1983); furthermore, in the extreme case where there are only two guides, denoted *g*_1_ and *g*_2_, the log-odds-ratios are perfectly anticorrelated 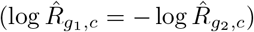, which straight-forwardly violates the PRDS definition (because 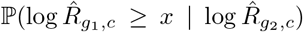 is not nondecreasing in 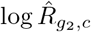). Therefore, the Benjamini-Hochberg procedure is not guaranteed to control the FDR. However, when the number of guides is large, the (anti)correlation of the multivariate hypergeometric components shrinks to 0, so assuming independence (which implies PRDS) may be a reasonable approximation.

Fishash therefore offers three options for controlling the FDR: in addition to Benjamini-Hochberg, we also provide options to use the Benjamini-Yekutieli procedure which controls FDR under arbitrary dependence (Benjamini and Yekutieli, 2001), as well as the procedure of Guo and Sarkar (2020) to control FDR assuming tests within a cell are dependent but tests across cells are independent. By default, fishash uses the Guo-Sarkar procedure, as a typical single-cell dataset contains (tens of) thousands of cells, so tests should be approximately independent across cells. However, if the number of cells is small, the Benjamini-Yekutieli procedure can be used to avoid assuming independence across cells; conversely, if the number of cells and guides are both large, the Benjamini-Hochberg procedure can used to increase power.

We briefly recount the multiple testing procedure of Guo and Sarkar (2020). For each cell, we define the minimum Bonferroni corrected p-value as

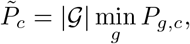

then we order the block p-values as 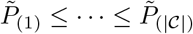 and find

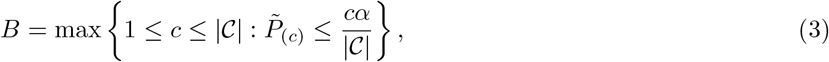

and finally reject 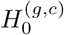 at significance level *α* if 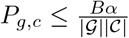, assuming *B* exists (if not, accept all null hypotheses).

Since 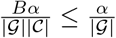, the procedure is bounded above by the within-cell Bonferonni correction, and therefore controls the FWER within each cell – the expected fraction of cells with an incorrect guide is at most *α*. In addition, Guo and Sarkar (2020) show that it controls the total FDR (across cells and guides) at level *α*.

### 2.4 Model for simulating gRNA counts

We adapted the Cellbender model (Fleming et al., 2023) to simulate guide RNA counts with a realistic model of background noise, while varying parameters such as signal-to-noise ratio (SNR), multiplicity-of-infection (MOI), and guide library size.

Cellbender models single-cell read counts as coming from three processes:

1. Signal: reads for molecules that were truly in the cell.
2. Endogenous noise: reads for molecules that were not in the cell but were present in its droplet, for example extracellular debris that wound up in the ambient solution.
3. Exogenous noise: reads that were not present in the cell nor its droplet, and are due to barcode swapping from sequencing or other artifacts, for example, chimeric reads due to PCR chimeras.

We recount the Cellbender hierarchical probabilistic model here. The count of feature *g* in cell *c* is written as the sum of the three processes,

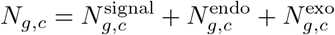

Where

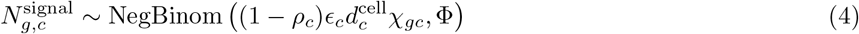

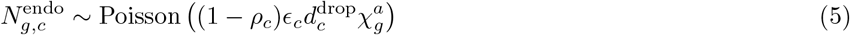

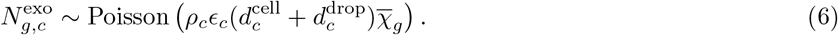

The physically encapsulated ambient molecules 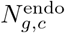 are generated from the endogenous ambient profile 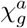, the droplet size factor 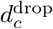, and the droplet-specific capture efficiency *ϵ*_*c*_. The barcode-swapped molecules 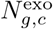 are generated from a droplet-specific rate *ρ*_*c*_ additionally modulated by the total physically captured molecules in the droplet 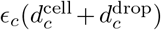 and the dataset-wide (pseudobulk) average expression 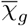. Finally, the true biological counts 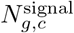 are modeled as a negative binomial with dispersion Φ and rate depending on the droplet-specific capture efficiency *ϵ*_*c*_, the non-chimeric fraction (1−*ρ*_*c*_), the cell size factor 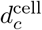, and the true latent expression of the cell *χ*_*g,c*_. Note that we follow the original Cellbender model (Fleming et al., 2023) in assuming the noise counts 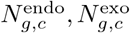 in (5), (6) follow a Poisson distribution conditional on the latent factors, but it is also possible to add additional overdispersion to these terms, which we consider in Appendix C.

A key difference of our simulation with the original Cellbender model is how we model the true latent expression *χ*_*g,c*_. Cellbender models *χ*_*g,c*_ as a neural network transformation of a lower dimensional Gaussian latent variable **z**_*g,c*_, similar to scVI (Lopez et al., 2018), which is a sensible model for gene expression but less so for guide RNA. Instead, the guide transcripts in Perturbseq data are much sparser, with at most a handful of lentiviral infections per cell.

A common experimental protocol is to infect cells at some low MOI, say 0.3, and then use a fluorescent marker to select cells with at least 1 guide. We therefore model the number of infections *K*_*c*_ as a Hurdle-Poisson model (i.e. zero-inflated or zero-deflated Poisson), where *λ*_MOI_ is the original Poisson rate and *θ* = ℙ (*K*_*c*_ = 0) is the hurdle probability due to imperfect selection. Conditional on the number of infections *K*_*c*_, we then sample the guides according to a multinomial distribution with probabilities ***π*** = (*π*_1_, …, *π*_|_𝒢_|_). If *J*_*g,c*_ the number of times *c* was infected by *g* is positive, then we assume *g* is expressed proportional to a guide-specific size factor 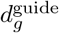. To summarize,

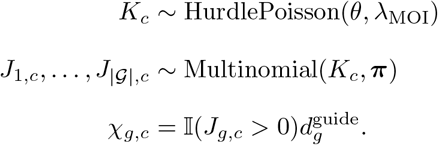

Unless otherwise stated, we use the following values for the variables above. We set the overdispersion of the signal counts Φ = 1. We set the droplet capture efficiency *ϵ*_*c*_ to be Gamma-distributed with mean 1 and shape 50 (the same default parameters used by Cellbender). We model the fraction of UMIs from exogenous noise (barcode swapping) per cell *ρ*_*c*_ as Beta distributed, with mean 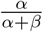 varying across simulations, and with the sum of its shape parameters *α* + *β* equal to 10. We model the cell size factors and ambient size factors 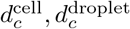 as log-normal with 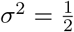 and their means varying across simulations (their means determine both the median number of UMIs per cell as well as the dataset’s signal-to-noise ratio SNR, which we vary across simulation settings). For the ambient background frequencies 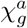, we model these as Dirichlet-distributed, with mean equal to abundance of guide *g* in the signal and total dispersion Σ_g_ *α*_*g*_ = |𝒢|. The motivation for having 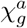 be centered around the true signal frequencies is because these are typically somewhat correlated with the ambient frequencies, due to ambient UMIs being formed from damaged cells. For the Hurdle-Poisson model, we typically set 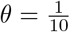 and 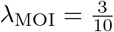, though we vary this in some simulations to study accuracy in different MOI regimes. We set the guide infection multinomial probabilities ***π*** ~ Dirichlet(1, …, 1) and the guide size factors as 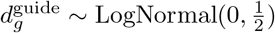.

### 2.5 Compared methods in benchmarks

For our benchmarks, the methods we compared against were crispat (Braunger and Velten, 2024), CLEANSER (Liu et al., 2025), DemuxEM (Gaublomme et al., 2019), geomux (Teyssier, 2024), and SCEPTRE (Barry et al., 2021). Unless otherwise stated below, all methods were run with default parameter values. We list the package versions used in Table 1.

**Table 1:** Software packages and versions used for benchmarking guide assignment methods.

| Package | Version |
| --- | --- |
| CLEANSER | 1.2 |
| Crispat | 0.9.10 |
| DemuxEM | 0.1.8 |
| Geomux | 0.5.5 |
| Fishash | 0.3.0 |
| SCEPTRE | 0.10.3 |

Crispat (Braunger and Velten, 2024) implements several mixture models for guide assignment; we included the Negative Binomial mixture, the Poisson mixture, the Poisson-Gaussian mixture (originally adapted from Replogle et al. (2022)), and the Gaussian mixture (which is similar to the model used by Cellranger-Multi). For the Gaussian mixture, we ran it in mode nonzero=True to only consider nonzero entries of the count matrix.

For SCEPTRE, we used the mixture method, which fits a latent variable Poisson GLM (Barry et al., 2024). Note this method requires matched gene expression data, as it includes the total gene expression count and features as covariates; since our simulation model doesn’t include gene expression, we used Splatter (Zappia et al., 2017) to generate fake expression data on the simulated data. Since the gene expression is uncorrelated with the guide counts in the simulated data, the SCEPTRE model would be approximately equivalent to a Poisson mixture model in this setting. Note the real data examples include matched gene expression data, so are a fairer comparison for SCEPTRE.

CLEANSER (Liu et al., 2025) models the guide counts as a mixture of signal and noise components, where the signal follows a Negative Binomial distribution, and the noise follows either a Poisson (“cropseq” mode) or Negative Binomial (“direct-capture” mode) distribution. We ran CLEANSER in both modes for all scenarios.

DemuxEM (Gaublomme et al., 2019) models the counts in each cell using a Dirichlet-Multinomial model, with an improper sparsity-inducing Dirichlet prior. It calls features as present when their posterior expected counts from signal are above some threshold, by default 10, but we lowered this to 2 due to the lower expression of gRNA data compared to HTOs (which DemuxEM was originally developed for).

Geomux (Teyssier, 2024) uses a hypergeometric model, similar to fishash. However, it does not include the block-dependent multiple testing correction (instead using the Benjamini-Hochberg procedure to control FDR (Benjamini and Hochberg, 1995)), nor the correction for Simpson’s paradox. Other differences are that it uses *N*_*g,c*_ − 1 instead of *N*_*g,c*_ in the hypergeometric test, which is similar to conditioning on *N*_*g,c*_ ≥ 1, and includes an adaptive procedure for setting a log-odds-ratio threshold.

We note that the methods were *not* all run with the same cutoffs, so may be expected to have different tradeoffs between precision and recall. Fishash and geomux were both run at 5% FDR cutoff, their default setting. We used a 80% posterior cutoff for csCLEANSER and a 50% cutoff for dcCLEANSER, which were the defaults in the CLEANSER paper (Liu et al., 2025). Crispat uses a hard-coded cutoff of 50% posterior probability, while we ran SCEPTRE with its default 80% probability cutoff. DemuxEM does not use a probability cutoff, but instead uses a cutoff based on the posterior mean for the number of reads coming from signal; by default it requires posterior expected signal count of at least 10, which typically works well for HTO data, but since gRNA data tend to have much lower counts, we lowered this to 2 (min signal=2) when running DemuxEM.

We call the cutoffs above the “default thresholds”. On a subset of datasets, we also explore different ranges of thresholds (Figures 5c, 6c; Appendix D). For fishash, geomux, CLEANSER, and SCEPTRE, we set the FDR or posterior cutoffs at {0.01, 0.05, 0.1, 0.2, 0.35, 0.5, 0.65, 0.8, 0.9, 0.95, 0.99}. For DemuxEM, we set its min signal cutoff at {1, 2, 3, 4, 5, 6, 7, 8, 9, 10, 15, 20}. For fishash and CLEANSER, we additionally generated “full” precision-recall (PR) curves based on the continuous test statistic (negative log p-value or posterior probability), and also included the full PR curve from the raw counts as a baseline. The other methods did not return test statistics for unassigned guides, so it was infeasible to compute full PR curves for them, and we only computed “discretized” PR curves based on their parameter sweeps. We did not generate PR curves for crispat because of its hard-coded posterior cutoff at 50%.

## 3 Results

We compare fishash against several other methods for assigning gRNAs, on simulated datasets with known ground truth, as well as real datasets with partial or proxy ground truth (Liu et al., 2025; Replogle et al., 2022).

### 3.1 Results on barnyard dataset

To benchmark guide assignment, Liu et al. (2025) generated a barnyard dataset consisting of mixed human and mouse cells, where the cells were infected with species-specific gRNA libraries. The human-mouse mixtures were prepared with two approaches, “non-co-cultured” and “72-h co-cultured”. For each mixture, CROP-seq and direct capture libraries were generated. For each sample, we filtered cells using the same gene expression QC metrics as in the original paper (requiring mitochondrial fraction *<* 15%, number of features between 1500 and 6000, number of UMIs between 3500 and 20000, and 90% of UMIs to be from a single species).

While the ground truth of which cell received which guide is unknown, a partial ground truth is available – human cells should only contain guides from the human library, while mouse cells should only contain guides from the mouse library. We therefore measured the accuracy of gRNA-based species assignment by computing the fraction of cells containing at least 1 guide from the correct species and 0 guides from the incorrect species.

We show the results in Figure 3 and Table 2. On the 2 Cropseq samples, the two hypergeometric-based approaches (geomux and fishash) performed best, with geomux having slightly higher accuracy but within 1 standard error of fishash. On the 2 direct capture samples, fishash was the most accurate method, but geomux was no longer as accurate, which was due to geomux not assigning any guides in a large number of cells (Figure 3a).

**Table 2:** Accuracy (with standard errors) of gRNA-based species assignment on the barnyard dataset of Liu et al. (2025).

|  | Cropseq mix0hr | Cropseq mix72hr | DirectCapture mix0hr | DirectCapture mix72hr |
| --- | --- | --- | --- | --- |
| cleanser_cs | $0.9414 \pm 0.0030$ | $0.9410 \pm 0.0027$ | $0.8612 \pm 0.0039$ | $0.9221 \pm 0.0028$ |
| cleanser_dc | $0.9178 \pm 0.0034$ | $0.8459 \pm 0.0041$ | $0.8614 \pm 0.0039$ | $0.9308 \pm 0.0027$ |
| crispat_gauss | $0.9367 \pm 0.0031$ | $0.9346 \pm 0.0029$ | $0.8200 \pm 0.0041$ | $0.8945 \pm 0.0030$ |
| crispat_negbinom | $0.4556 \pm 0.0007$ | $0.4815 \pm 0.0040$ | $0.8865 \pm 0.0037$ | $0.9381 \pm 0.0025$ |
| crispat_poisgauss | $0.9501 \pm 0.0028$ | $0.9502 \pm 0.0025$ | $0.8739 \pm 0.0039$ | $0.9331 \pm 0.0026$ |
| crispat_poisson | $0.9086 \pm 0.0036$ | $0.9169 \pm 0.0032$ | $0.8763 \pm 0.0041$ | $0.8956 \pm 0.0035$ |
| demuxem | $0.9364 \pm 0.0031$ | $0.9364 \pm 0.0028$ | $0.8975 \pm 0.0036$ | $0.9408 \pm 0.0025$ |
| fishash | $0.9563 \pm 0.0026$ | $0.9533 \pm 0.0025$ | **$0.9100 \pm 0.0034$** | **$0.9491 \pm 0.0024$** |
| geomux | **$0.9578 \pm 0.0025$** | **$0.9555 \pm 0.0024$** | $0.7471 \pm 0.0044$ | $0.5252 \pm 0.0056$ |
| sceptre_mixture | $0.9491 \pm 0.0028$ | $0.9472 \pm 0.0026$ | $0.8953 \pm 0.0036$ | $0.9408 \pm 0.0025$ |

**Figure 3:**
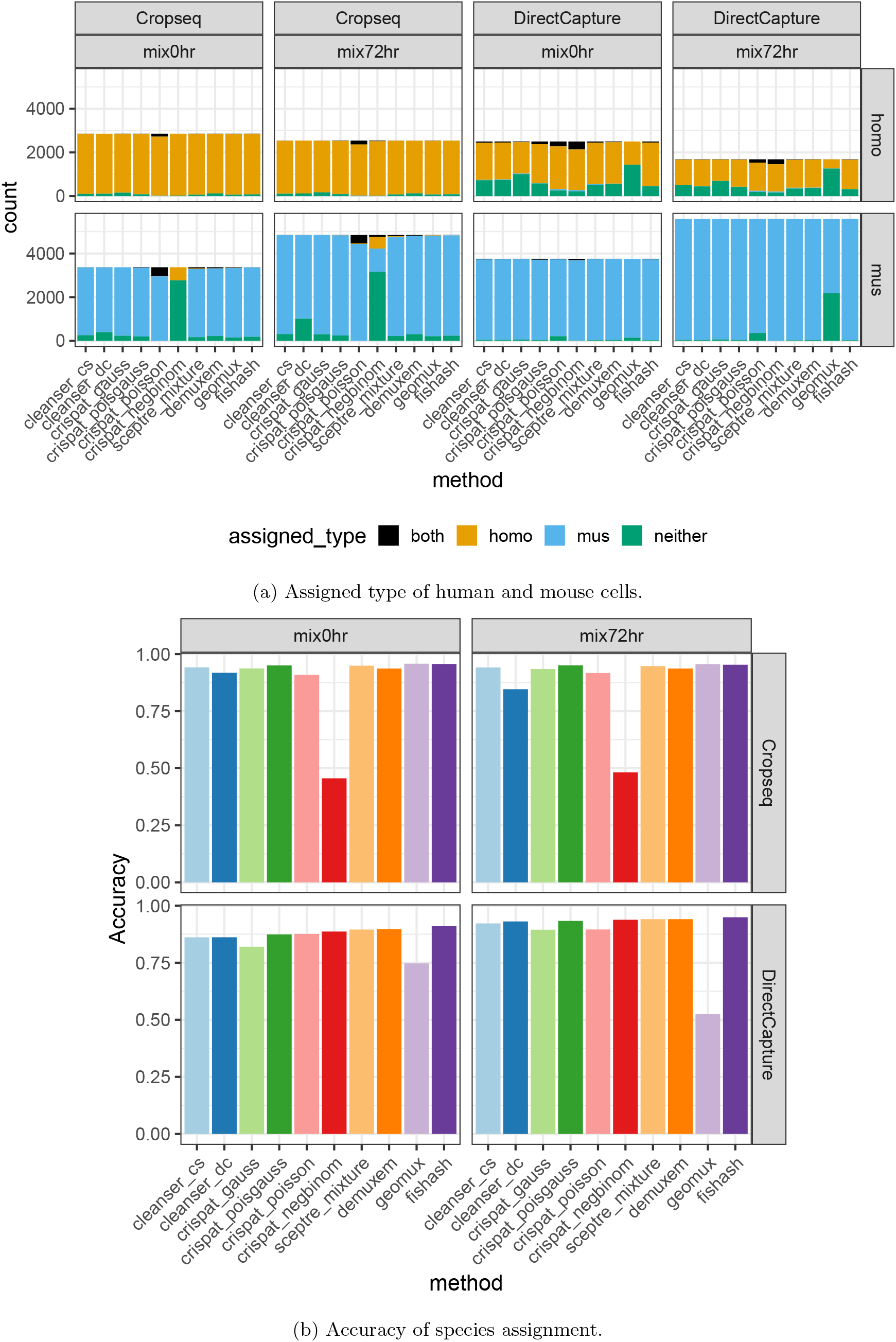
Performance of guide assignment methods on the barnyard dataset of Liu et al. (2025). This dataset includes a mixture of human and mouse cells, which were infected with separate guide libraries. Figure 3a shows whether each cell was assigned human guides, mouse guides, both, or neither. Figure 3b plots the accuracy of the species assignment, which we define as the fraction of cells with at least 1 guide from the correct species and 0 guides from the incorrect species.

### 3.2 Results on K562 genome-wide perturbseq

In a landmark study, Replogle et al. (2022) performed genome-wide perturbseq on a K562 cell line. While this dataset does not contain a known ground truth, the study uses a dual-guide CRISPRi lentiviral construct, and we propose to construct a proxy ground truth based on the concordance of the paired guides. Specifically, we labelled cells as truly containing a guide if it had at least 1 read for both the guide and its matched pair. The logic for this proxy ground truth is that it is unlikely for ambient or chimeric reads from both guides of a perturbation to co-occur in a droplet by chance, since there are a total of 22,588 sgRNA targets (11,294 sgRNA pairs) in the dataset, while the median droplet contains reads for 9 sgRNAs. Crucially, note that each guide has a separate promoter, and thus each of the 2 guides are transcribed onto physically distinct transcripts.

We plot F1, precision, and recall metrics based on this proxy ground truth in Figure 4. Among the benchmarked methods, Fishash had the highest F1 score in 78% (213 / 273) of 10X runs. Additionally, Fishash had the highest recall among methods (median 97.1%; the medians for other methods ranged from 60.1-95.4%), while having a comparable level of precision to other methods (median 68.7%; the medians for other methods ranged from 62.8-70.7%). We also included a baseline of calling all guides with at least 1 read (with a median precision of 14.8%; note by definition of the proxy ground truth, this baseline has a recall of 100%).

**Figure 4:**
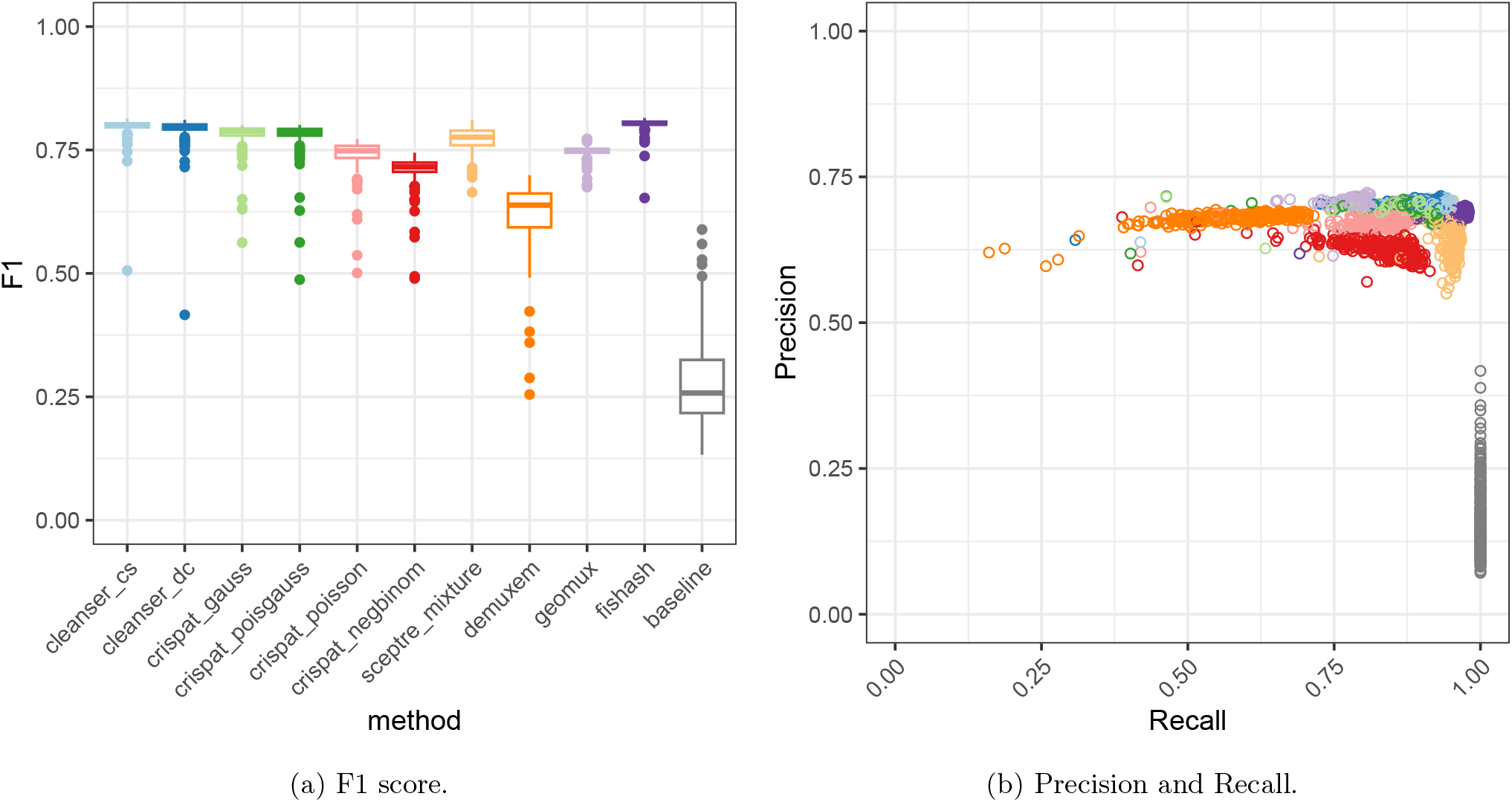
Accuracy metrics on the K562 genome-wide perturbseq (Replogle et al., 2022), based on a proxy ground truth constructed from the dual-guide system (we labelled cells as truly containing a guide if it contained reads for both the sgRNA and its paired guide). We included a baseline which consists of assigning all guides with at least 1 read. Fishash had the highest F1 score in 78% (213/273) of batches.

**Figure 5:**
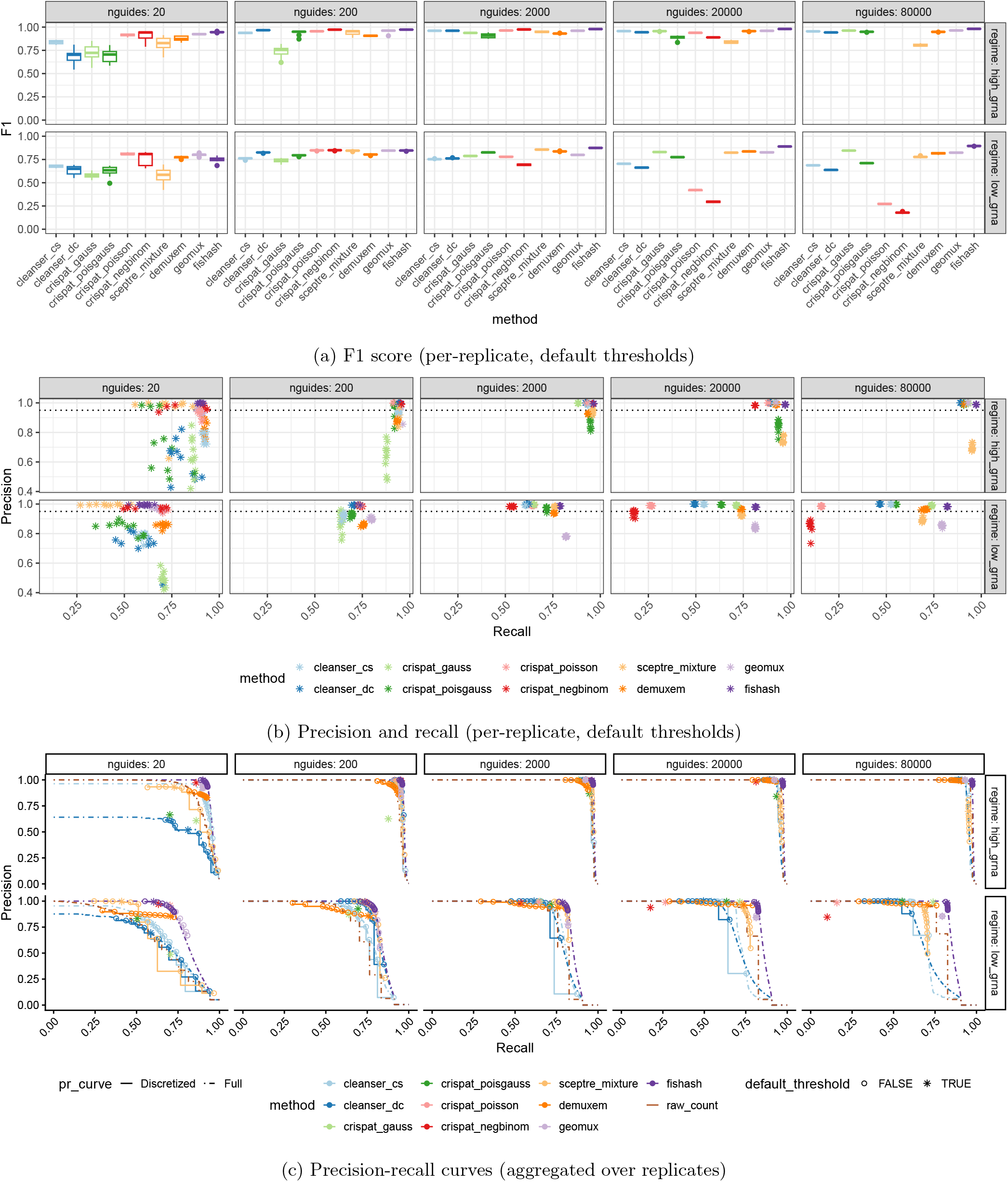
Guide calling performance as the the number of guides varies, in two regimes (high-gRNA and low-gRNA expression). Each scenario had 10 replicates with 20,000 cells each. Some crispat models were not included in the largest setting (80,000 guides) due to long running time (over 72 hours). In 5b, the dotted line at 0.95 precision is the nominal cutoff for fishash and geomux (5% FDR). In 5c, precision-recall curves were computed using discrete parameter sweeps (“Discretized”) and/or over a full range (“Full”), depending whether the method had tunable arguments and/or returned continuous test statistics (methods with neither are plotted as a single point).

We note that no method achieves a precision above 71%, despite many of methods using a higher nominal precision cutoff. Specifically, fishash and geomux used FDR cutoffs of 5%, corresponding to a nominal precision of 95%, while csCLEANSER and SCEPTRE used posterior cutoffs of 80%. One possible cause for the loss of precision is epigenetic silencing of one of the promoters, leading to only one of the guides being expressed and causing the proxy ground truth to mistakenly label such cells as negative.

### 3.3 Results on simulated datasets

We next use the simulation framework in Section 2.4 to benchmark the performance of guide assignment methods across a range of parameters including signal-to-noise ratio (SNR), multiplicity-of-infection (MOI), and size of guide library. All simulations consisted of 20,000 cells.

We first examined how varying the number of unique guides in the library affected performance. We varied the number of unique guide barcodes between 20, 200, 2000, 20,000, and 80,000. We considered two regimes:

1. A high-gRNA expression regime, with a median of 100 UMIs per cell, SNR of 4, and with 25% of the noise to be exogenous (due to barcode swapping rather than being present in the droplet).
2. A low-gRNA expression regime, with a median of 20 UMIs, SNR of 1, and with 75% of the noise to be exogenous. The higher level of exogenous noise in this scenario is motivated by the fact that the experimentalist may have to perform more PCR cycles to amplify sufficient material in the low-expression regime, leading to higher levels of barcode swapping.

Both regimes used an MOI of 0.3, and a hurdle probability (chance of non-infection) at 0.1.

Note that jobs were submitted on a slurm queue with a 72 hour time limit, and models that failed to complete all jobs for a scenario within the alloted time were excluded. Specifically, the crispat Poisson and Negative Binomial mixtures were excluded from the high-expression 80,000 guide scenario.

We show the results of this experiment in Figure 5. Of the 100 simulations (10 replicates each from 10 scenarios), fishash had the highest F1 score in 70 of 100 simulations, and the highest median F1 score in 8 of 10 scenarios. Additionally, fishash successfully controlled the FDR at the nominal level (precision ≥ .95) in all 100 simulations. By contrast, geomux failed to control FDR at its nominal 5% level, especially in the low gRNA expression regime; CLEANSER and SCEPTRE also sometimes failed to control precision at the nominal level when varying their posterior cutoff (Figure D.3a). When varying the assignment threshold, fishash had the highest AUPRCs in several settings, especially when the number of guides was large (Figure 6c, Figure D.1).

**Figure 6:**
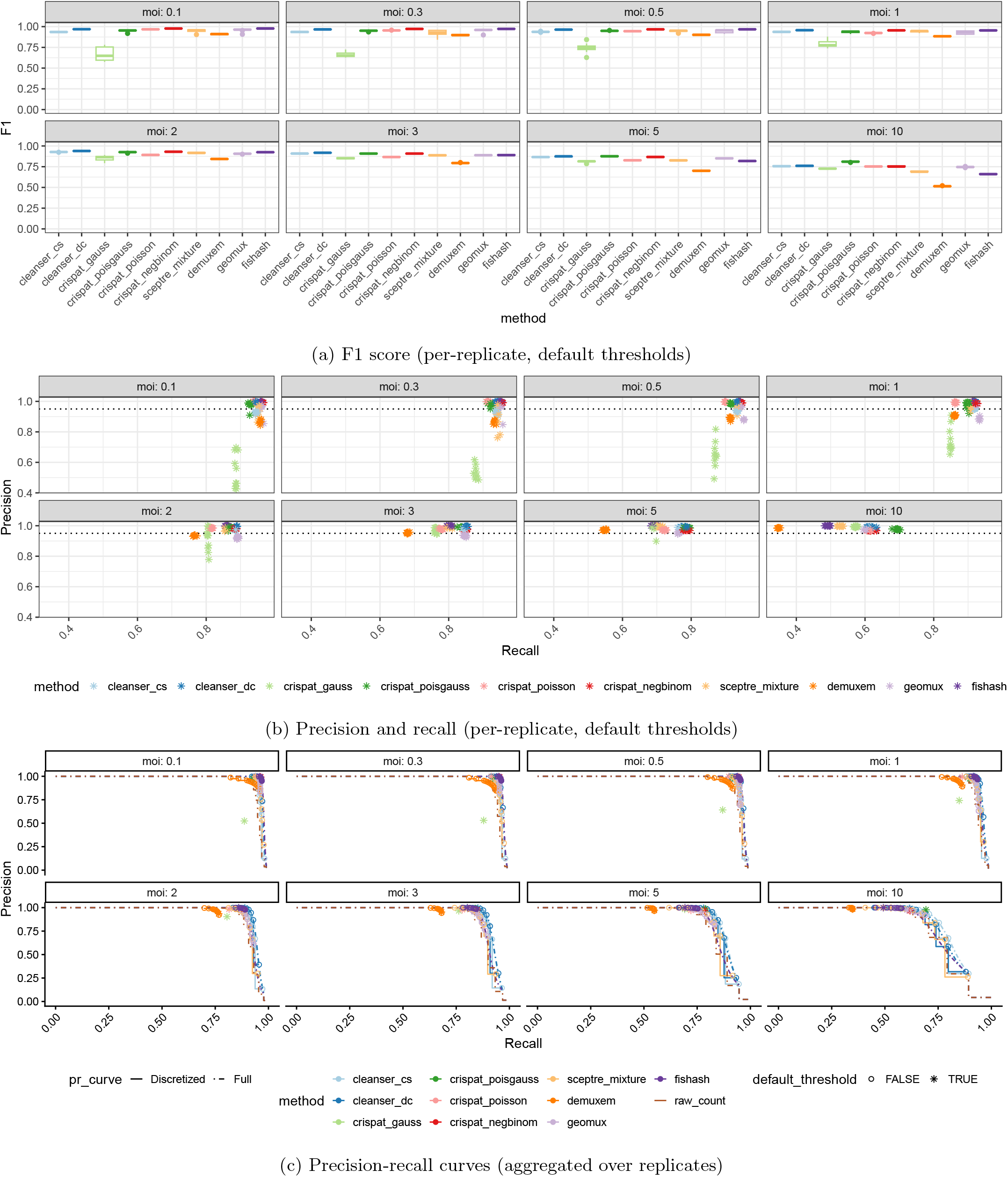
Precision, recall, and F1 scores while varying the multiplicity-of-infection (MOI) in simulations. The number of guides was fixed at 200. Aside from MOI and number of guides, most parameters were the same as in the high-gRNA simulation scenario of Figure 5.

Next, we examined how varying the MOI affects performance. We simulated 20,000 cells with 200 guides and MOI varying between 0.1, 0.3, 0.5, 1, 2, 3, 5, and 10. Other simulation parameters were set at at the same values of the high-gRNA expression scenario (100 UMIs per cell, SNR of 4, 25% of noise is exogenous, hurdle probability 0.1). We show the results in Figure 6. Fishash has the highest median F1 score for the low MOI scenarios (MOI ≤ 0.5), while dcCLEANSER had the highest median F1 when 1 ≤ MOI ≤ 3, and crispat Poisson-Gaussian mixture had the highest median F1 when MOI ≥ 5. While fishash did not perform as well on the high MOI scenarios according to the F1 score, when considering the full (exact) AUPRC its gap with other methods was smaller (Figure 6c, Figure D.1). Therefore, in these cases it may be beneficial for fishash to manually select an alternative threshold based on visualization of the test statistic (Figure D.4).

We also consider the case of highly-overdispersed noise counts in Appendix C. There, we modify the above simulation scenarios to allow for extreme overdispersion of the noise counts 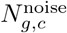, by assuming they are conditionally Geometric instead of Poisson. In that case, fishash no longer controls the FDR at the nominal level in all settings; however fishash continues to have the highest F1 score in several settings, especially when the number of guides is large.

Finally, we examined in what contexts the correction for Simpson’s paradox has a noticeable effect on fishash. Specifically, note that Simpson’s paradox does not occur when the signal guide frequencies equal the noise guide frequencies (Proposition 1). Therefore we simulated three regimes of correlation between the signal guide frequencies and the endogenous noise frequencies 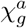, showing the results in Figure 7. Specifically, the three correlation regimes were:

**Figure 7:**
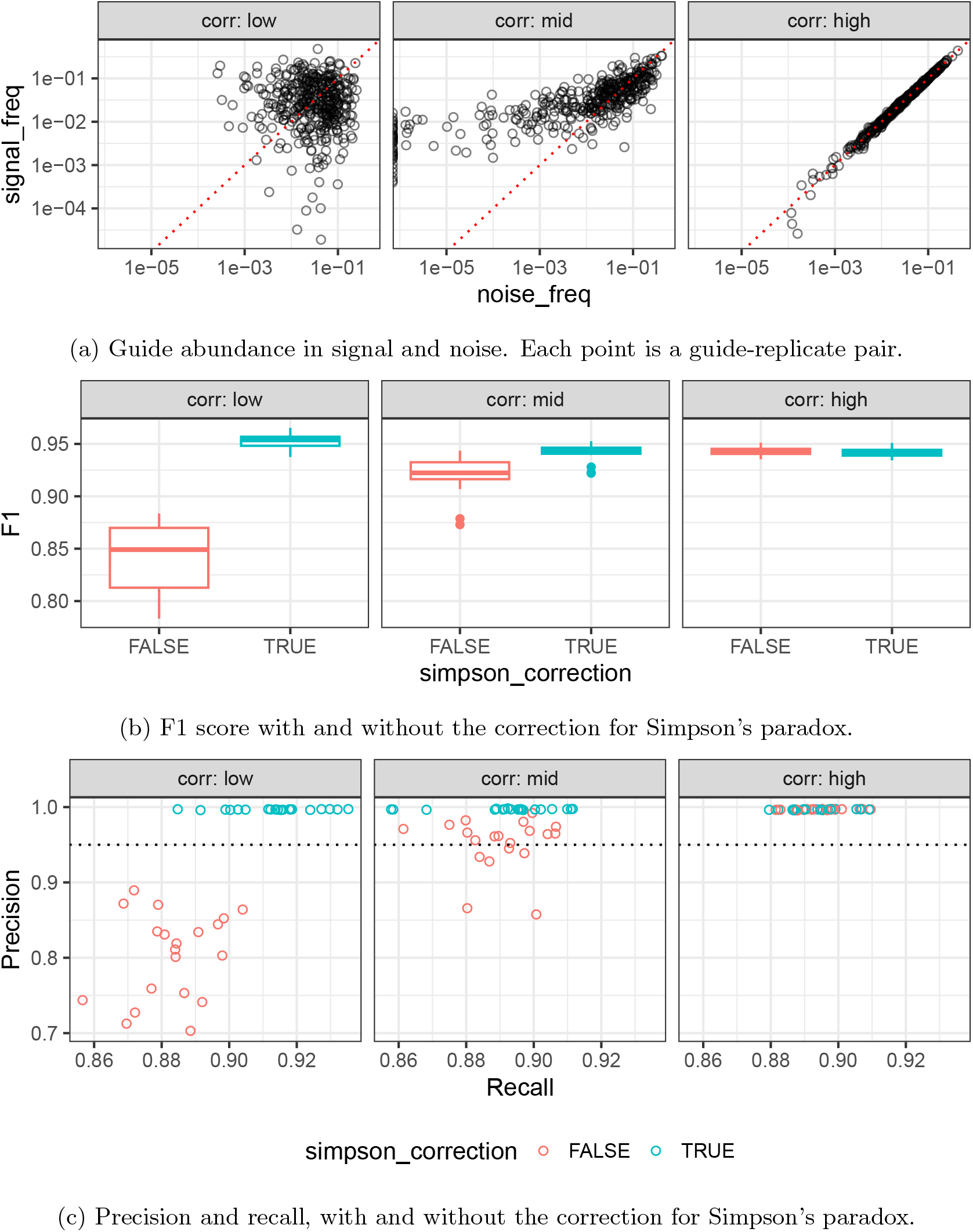
Effect of Simpson’s paradox correction in three regimes for the correlation between the total reads a guide has among signal and noise reads. In the low correlation regime, the correction yields a substantial improvement in both precision and recall, and the uncorrected Fisher test fails to control FDR at the nominal level (5%).

1. Low correlation regime: 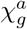 is sampled from a uniform distribution, i.e. a Dirichlet distribution with shape parameters (*α*_1_, …, *α*_|_𝒢_|_) = (1, …, 1).
2. Medium correlation regime: 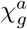 is sampled from a Dirichlet distribution centered around the signal frequencies, with the sum of the shape parameters given by Σ_*g*_ *α*_*g*_ = |𝒢|.
3. High correlation regime: 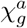 equals the signal frequencies.

We set the exogenous noise to 0 in all three settings, and simulated 20 replicates per scenario with 20 guides each. Other parameters were the same as in the high-expression regime: we used a median of 100 UMIs per cell, SNR of 4, MOI of 0.3, and hurdle probability of 0.1.

In Figure 7, we visualize the relationship between the signal and noise guide frequencies in each regime, as well as the F1 score, precision, and recall of fishash with and without the correction for Simpson’s paradox. In the low correlation regime, the correction yields a substantial improvement in precision and recall, and the uncorrected Fisher test fails to control the FDR at the nominal level (5%) in all replicates. In the medium correlation regime, the correction for Simpson’s paradox yields modest but noticeable improvements in F1 and precision, with the uncorrected Fisher test failing to control FDR in 6/20 simulations. In the high correlation regime, the performance with and without correction is negligible, with the uncorrected Fisher test having a slightly higher F1 score (.9426 vs .9421).

### 3.4 Running time and memory usage

In Figure 8, we show the resource usage as reported by Nextflow (Di Tommaso et al., 2017) for the simulation scenario of Figure 5, varying the guide library size from 20 to 80,000. All methods were run with 8 CPUs, except for fishash and geomux which are not parellelized and were allocated a single CPU. As noted previously, some crispat models were not included in the largest setting (80,000 guides) due to their long running time. An important caveat is that the jobs were run on a large HPC cluster via Slurm, so these metrics only give a rough idea of runtimes rather than a precise benchmark, because the jobs were run across multiple compute nodes which may have different performance characteristics (e.g. due to hardware, load, or node health).

**Figure 8:**
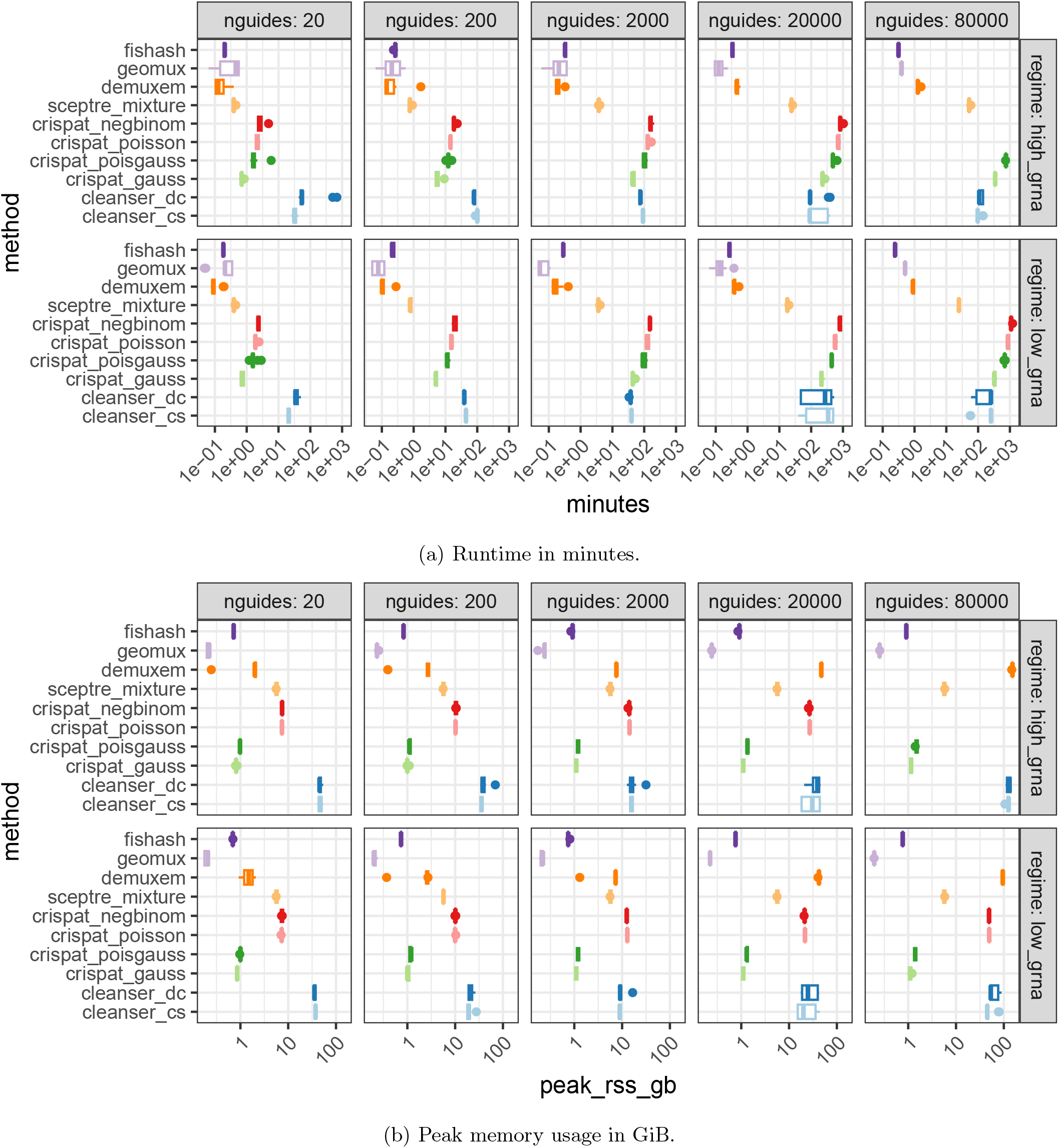
Resource usage (as reported by Nextflow) on the simulation scenario with varying guide library size from 20 to 80,000. Some crispat models were not included in the largest scenario (80,000 guides) due to their long running times. We note that the runtimes should only be used as rough estimates for performance rather than a precise benchmark, since the jobs were run across multiple compute nodes in a large, heterogenous HPC cluster via Slurm.

Fishash and geomux had the best resource usage overall, and could be run interactively on a laptop even in the largest scenarios (with runtimes in seconds and *<* 1 GiB memory usage). SCEPTRE had moderate memory usage (up to 6 GiB) and runtimes (up to 1 hour) and could also be run on a laptop, though less comfortably than fishash and geomux. DemuxEM had similarly fast runtimes as fishash and geomux, but required high amounts of memory (over 100 GiB) in the largest scenarios. Conversly, the crispat Gaussian mixture and Poisson-Gaussian mixture had low memory usage in all scenarios (≈ 1 GiB) but took several hours on larger scenarios. The crispat Poisson and Negative Binomial models had heavier memory and runtime requirements, and were excluded from the high-expression 80,000-guide scenario due to long running times (over 72 hours). Compared to the crispat Poisson and Negative Binomial models, CLEANSER had higher memory usage in the low-guide scenarios but similar memory usage in the high-guide scenarios, while having slower runtimes in the low-guide scenarios but faster runtimes in the high-guide scenarios (though still taking several hours in the latter).

## 4 Discussion

A benefit of the low computational cost of fishash is that it enables the analyst to assign guides during exploratory data analysis. While in many cases guide assignment can be run as a non-interactive pipeline step, some datasets may benefit from an interactive approach – for example, if a dataset contains multiple celltypes with different levels of guide expression, it may be beneficial to process the different celltypes separately. Exploratory analysis may also be beneficial to inspect how adjusting the FDR cutoff affects downstream results.

Potential methodological improvements could be made to both the multiple testing correction procedure and the correction for Simpson’s paradox. In particular, the multiple testing correction of Guo and Sarkar (2020) can be made more powerful by incorporating an adaptive estimate of the number of true positives. Additionally, the binary mask Ω_*g,c*_ used in the correction for Simpson’s paradox could be relaxed to allow for continuous values in [0, 1]; such a relaxed procedure may correspond to a Bayesian interpretation of Ω_*g,c*_ as the probability of coming from signal or noise.

In addition to our guide assignment method, we anticipate that our simulation framework will be useful for further development and benchmarking of guide assignment methods.

An interesting result of our simulations is that the the performance of guide assignment methods may vary considerably across dataset parameters such as SNR, MOI, and guide library size. This has some implications for experimental design – a common practice when running large screens is to run smaller pilot screens and develop pipelines on the smaller pilots, but our results suggest that performance of methods can change as the guide library size increases, which should be kept in mind when scaling up screens.

No single guide assignment was uniformly the best across all simulations and scenarios, however fishash was the top performing method in many cases, and performed particularly well when the number of guides was large. Its strong performance on large guide libraries, along with its speed, makes fishash particularly well suited for genome-scale perturbseq screens.

## 5 Conclusion

Compared to more sophisticated and computationally expensive approaches based on mixture models, using Fisher’s test to assign guides in Perturb-seq data performs surprisingly well. However, in order to control FWER and FDR at specified rates, it is important to supplement Fisher’s test with a multiple testing approach that accounts for the correlation between tests, as well as to correct for hidden confounding that causes Simpson’s paradox. We combine these elements to form a guide assignment algorithm that is highly performant in both speed and accuracy. We provide our method in an easy to use R package, fishash.

## Declarations

### Ethics approval and consent to participate

Not applicable.

### Consent for publication

Not applicable.

### Availability of data and materials

The Liu et al. (2025) barnyard dataset re-analyzed in this study is available at https://www.ncbi.nlm.nih.gov/geo/query/acc.cgi?acc=GSE272457. The Replogle et al. (2022) K562 genome-wide perturbseq dataset re-analyzed in this study is available at https://plus.figshare.com/articles/dataset/_Mapping_information-rich_genotype-phenotype_landscapes_with_genome-scale_Perturb-seq_Replogle_et_al_2022_MTX_files/20127869/1. The benchmark-ing results can be generated from the scripts available at https://github.com/jackkamm/fishash_analysis.

### Authors’ contributions

JK co-conceived the statistical method and study design, wrote the software, performed data analysis, wrote the initial manuscript draft, and created figures. JY co-conceived the statistical method and simulation study, performed software testing, and created figures. WF co-conceived the statistical method, substantively revised the manuscript, and supervised the work.

## Acknowledgments

We thank Oleg Mayba for extensive feedback that improved this manuscript. We also thank Jorge Kageyama, Bo Li, Yiming Yang, Stefan Peidli, Ana Meireles, and the Roche/Genentech AI Biology and Translation (AIBT) department for useful discussions and feedback during this work. We also thank the anonymous reviewers for helpful suggestions to improve the paper.

## A Proof of Proposition 1

Assume that guide *g* is not present in cell *c*, so 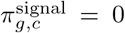. Furthermore, assume that barcodes *g* and *c* are independent in the noise reads, so that we can write 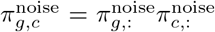 and 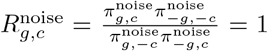. We prove that *R*_*g,c*_ *>* 1 if and only if 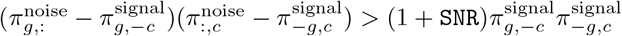.

We can expand *R*_*g,c*_ as

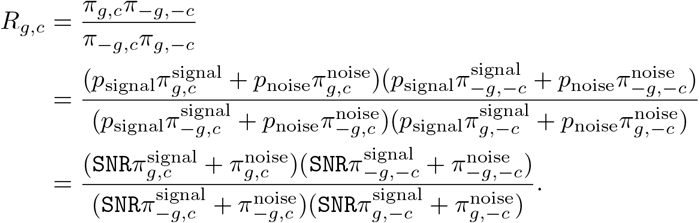

Since 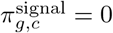, we have that *R*_*g,c*_ *>* 1 iff

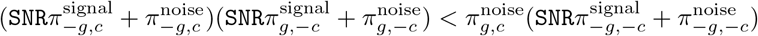

Since 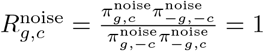, we can subtract 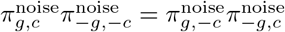 from both sides to obtain

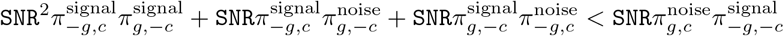

Dividing both sides by SNR and rearranging all terms to the RHS, this becomes

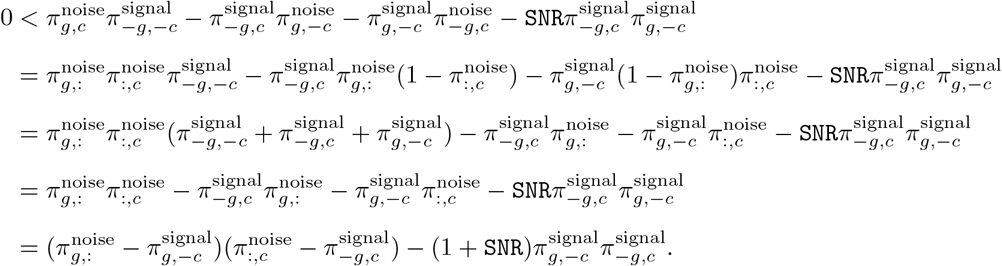

Finally, note that 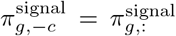 and 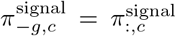, since by assumption 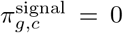. Substituting and rearranging terms yields

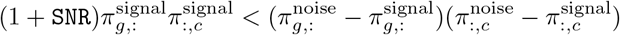

which completes the proof.

## B Details for Simpson’s Paradox correction

### B.1 Motivation for adjusted odds ratio 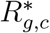

To correct for Simpson’s Paradox, we propose to test an adjusted odds ratio 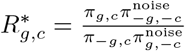, with null hypothesis 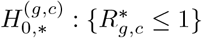 and alternative 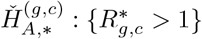. To motivate this correction, note that when 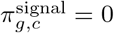,

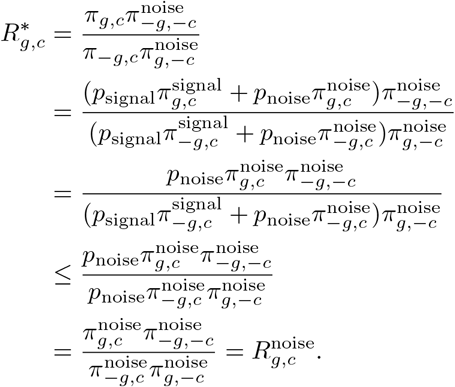

Note that 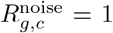 if guide barcode *g* and cell barcode *c* are independent among the noise counts, for example if the noise reads are sampled from the same Multinomial or Poisson distribution for all cells. Therefore, we have that 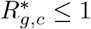 whenever guide *g* is not present in cell *c* and the barcodes *g, c* are independent in the noise UMIs.

### B.2 Estimating noise counts with rank-1 Poisson completion

If the terms 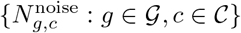 were observed, then we could test 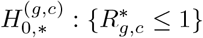 using Fisher’s exact test, with p-value 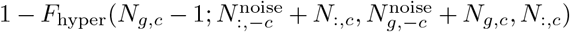. However, 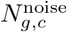are not observed; therefore, we propose to construct an approximate p-value *P*_*g,c*_ based on estimated noise counts,

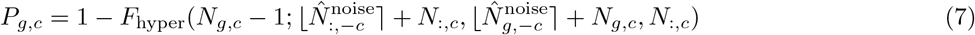

where the 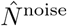 are plug-in estimators for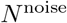, and ⌈*x*⌉ denotes rounding *x* to the nearest integer.

To estimate 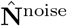, we take an iterative approach, first computing p-values *P*_*g,c*_ and assigning guides at FDR level *α*, then masking the assigned entries of **N** and imputing their values to estimate 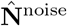, and then repeating.

In particular, let Ω ∈ {0, 1}^| 𝒢 |*×*| 𝒞 |^ denote a mask on 𝒢 ×𝒞, and let 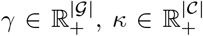 denote guide and cell abundances in the noise UMIs, respectively. Define *l*(*N*, Ω, *γ, κ*) as the Poisson log-likelihood of the masked counts given *γ, κ*,

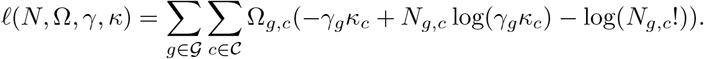

Let 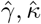 be the MLE estimates

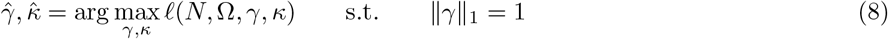

where the constraint ∥*γ*∥_1_ = 1 is added for identifiability. We then define 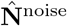 as

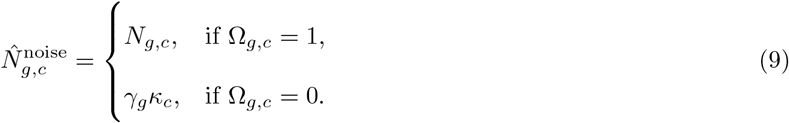

In other words, 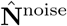 is the rank-1 matrix completion of **N** with mask Ω and Poisson loss function. To compute 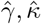, we alternate between optimizing *κ* for fixed *γ* and vice versa. In particular, for fixed *γ*, the optimal *κ*_*c*_ can be computed as 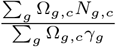. Note this approach is similar in flavor to alternating least squares solutions for low-rank matrix completion (Hastie et al., 2015), but substituting the squared-error-loss with the negative Poisson log-likelihood.

To set the mask Ω^(*i*)^ at iteration *i*, let 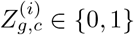 denote whether (*g, c*) was assigned at the current iteration,

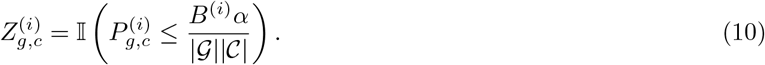

We then set

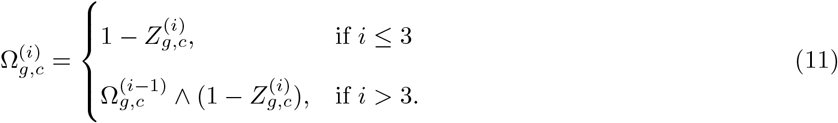

That is, entries that were assigned in the current iteration are masked; and in addition, for higher iterations (*i >* 3), entries that were masked in the previous iteration remain masked. The motivation for masking entries from earlier iterations is because such entries are often borderline-significant, so worth masking, and also may flip between significant or not in alternating iterations, preventing the algorithm from converging if we do not keep them masked. However, the p-values in the initial iterations can change more substantially, so we recompute the mask anew in the initial iterations (*i* ≤ 3).

## C Effect of additional noise overdispersion in simulations

Here we consider the effect of adding additional overdispersion to the noise counts in the simulations. Specifically, we modify the terms 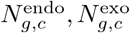 defined in (5) and (6) in Section 2.4, so that

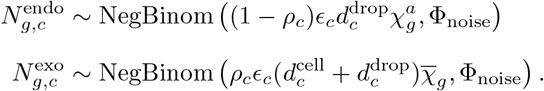

The case Φ_noise_ = 0 corresponds to the original definitions in (5) and (6), which in turn follow the original Cellbender model (Fleming et al., 2023). We note that even when Φ_noise_ = 0, the noise counts are overdispersed 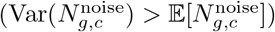 due to the variance of the latent terms *ρ*_*c*_, *ϵ*_*c*_,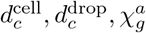. However, conditional on these latent variables, the noise counts 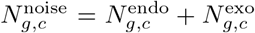 follow a rank-2 Poisson structure. The assumption that the noise counts follow a conditional low-rank Poisson or Multinomial structure is common in models for single-cell RNAseq contamination, including Cellbender (Fleming et al., 2023), SoupX (Young and Behjati, 2020), DecontX (Yang et al., 2020), and scAR (Sheng et al., 2022).

Nevertheless, there may be additional unmodeled sources of variation; therefore, we generated supplementary simulations to consider extra overdispersion on the noise counts. Specifically, we regenerated the varying-guide scenario of Figure 5 and the varying-MOI scenario of Figure 6, but added extra overdispersion to the noise counts by setting Φ_noise_ = Φ = 1.

We show the results in Figures C.1 and C.2. In terms of the F1 scores, the performance of fishash is qualitatively similar to the original results (Figure 5, 6): in the varying-guides scenario (MOI 0.3), fishash has the top median F1 score in 6/10 settings, and does particularly well in settings with high number of guides and/or low gRNA expression; while in the varying-MOI scenario (200 guides), fishash has relatively low F1 in the high-MOI regime while remaining close to the top in the low-MOI regime (though it no longer has the top median F1 score in any scenario, being supplanted by crispat-NB and dcCLEANSER). Likewise, the recall metrics are roughly similar to Figures 5 and 6; however the precision scores for many methods, and especially fishash, drop noticeably in the high overdispersion regime. In particular, fishash no longer controls FDR at the nominal 5% level in all scenarios.

We note that the Φ_noise_ = 1 setting we used in this simulation corresponds to a very high level of overdispersion – in particular, it corresponds to a Geometric distribution, which has the highest level of overdispersion among all Negative Binomial distributions with an integer-valued parameter for the number of successes. It is therefore unsurprising that the precision should drop in this setting due to the fatter right tail of the noise counts. Never-theless, while Fishash no longer has the highest precision among other methods in this scenario, it still maintains a competitive tradeoff between precision and recall, as indicated by its F1 scores.

## D Additional plots relating to precision-recall curves

**Figure C.1.**
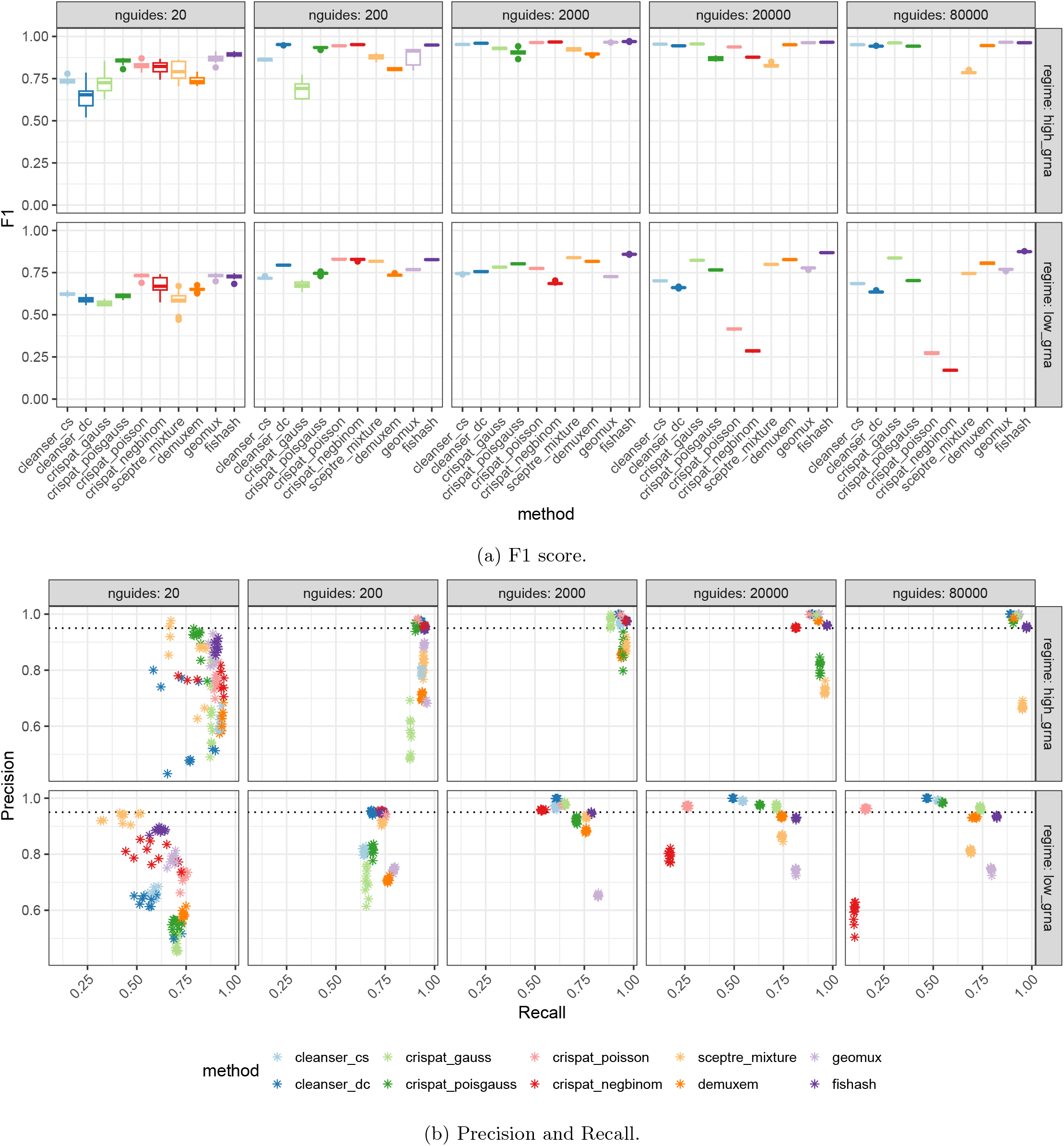
Guide calling performance when adding extra overdispersion to the simulation scenario of Figure 5, varying the number of guides in high-expression and low-expression regimes. The extra overdispersion was added by replacing the Poisson distribution with a Geometric distribution when simulating the noise counts.

**Figure C.2.**
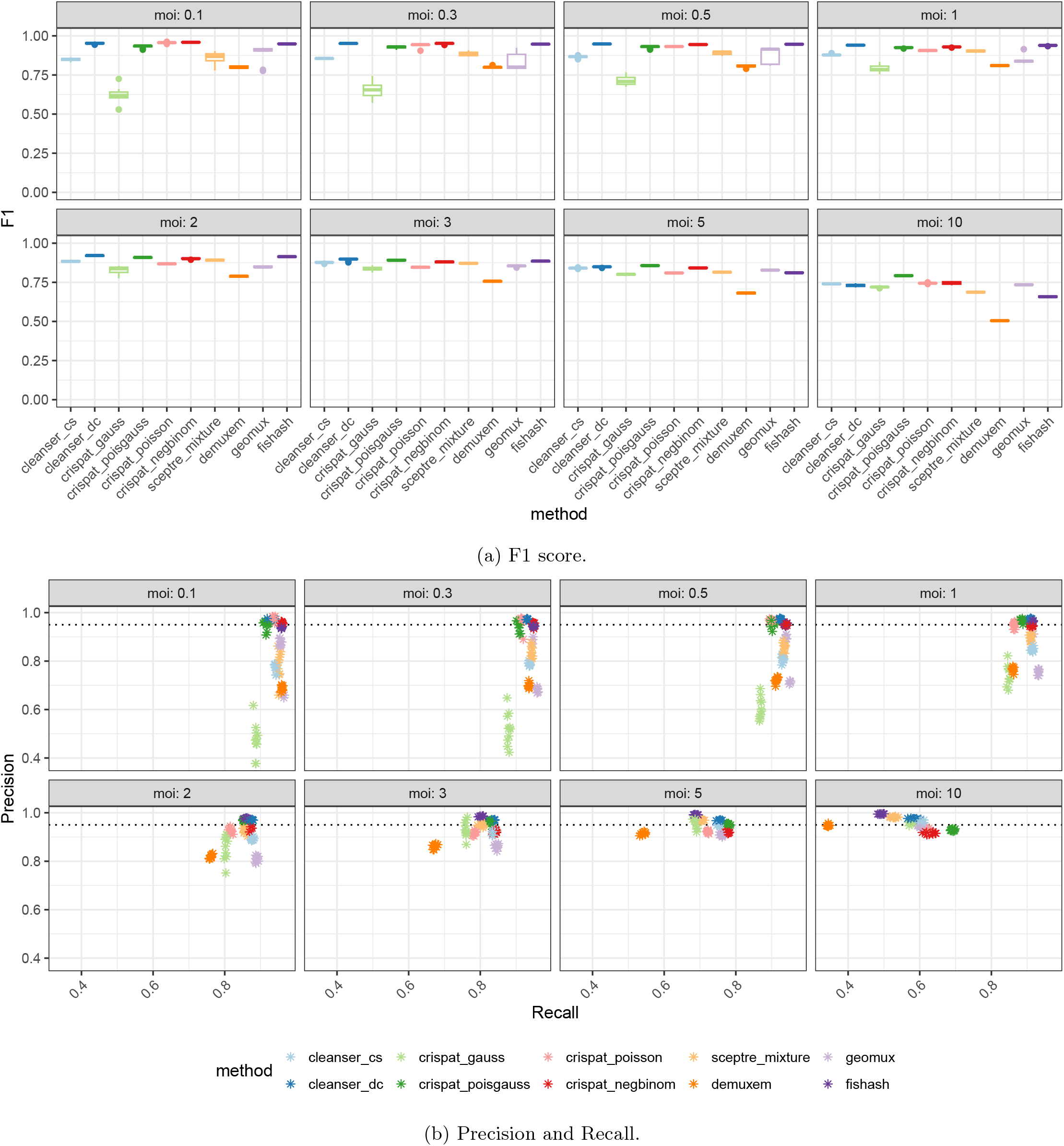
Guide calling performance when adding extra overdispersion to the simulation scenario of Figure 6, varying the MOI with a fixed number of guides (200). The extra overdispersion was added by replacing the Poisson distribution with a Geometric distribution when simulating the noise counts.

**Figure D.1.**
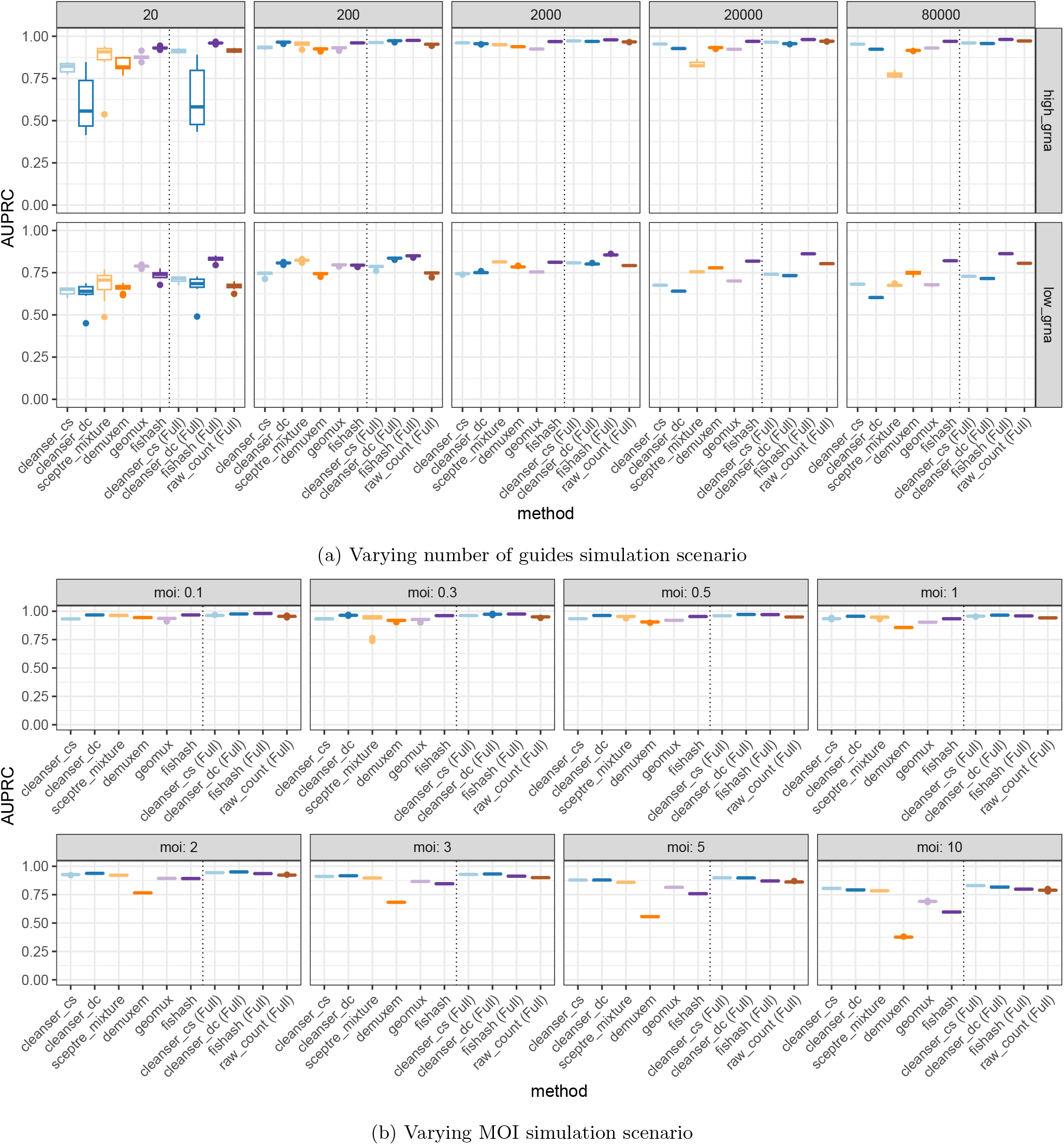
AUPRCs in the two main simulation scenarios. Discretized precision-recall curves were computed with a parameter sweep; additionally, full precision-recall curves were computed for methods that output continuous test statistics. The discretized AUPRC is generally an underestimate of the full AUPRC, especially when the discretized precision or recall values sweep a narrow subset of [0, 1] (cf Figures 5c, 6c). In the varying guide numbers scenario (D.1a), fishash has the top median AUPRC in 6 or 10 out of 10 settings, depending on if the discretized or full AUPRC is used. In the varying MOI scenario (D.1b), dcCLEANSER performed best, with the top median AUPRC in 6 (discretized) or 4 (full) settings out of 8.

**Figure D.2.**
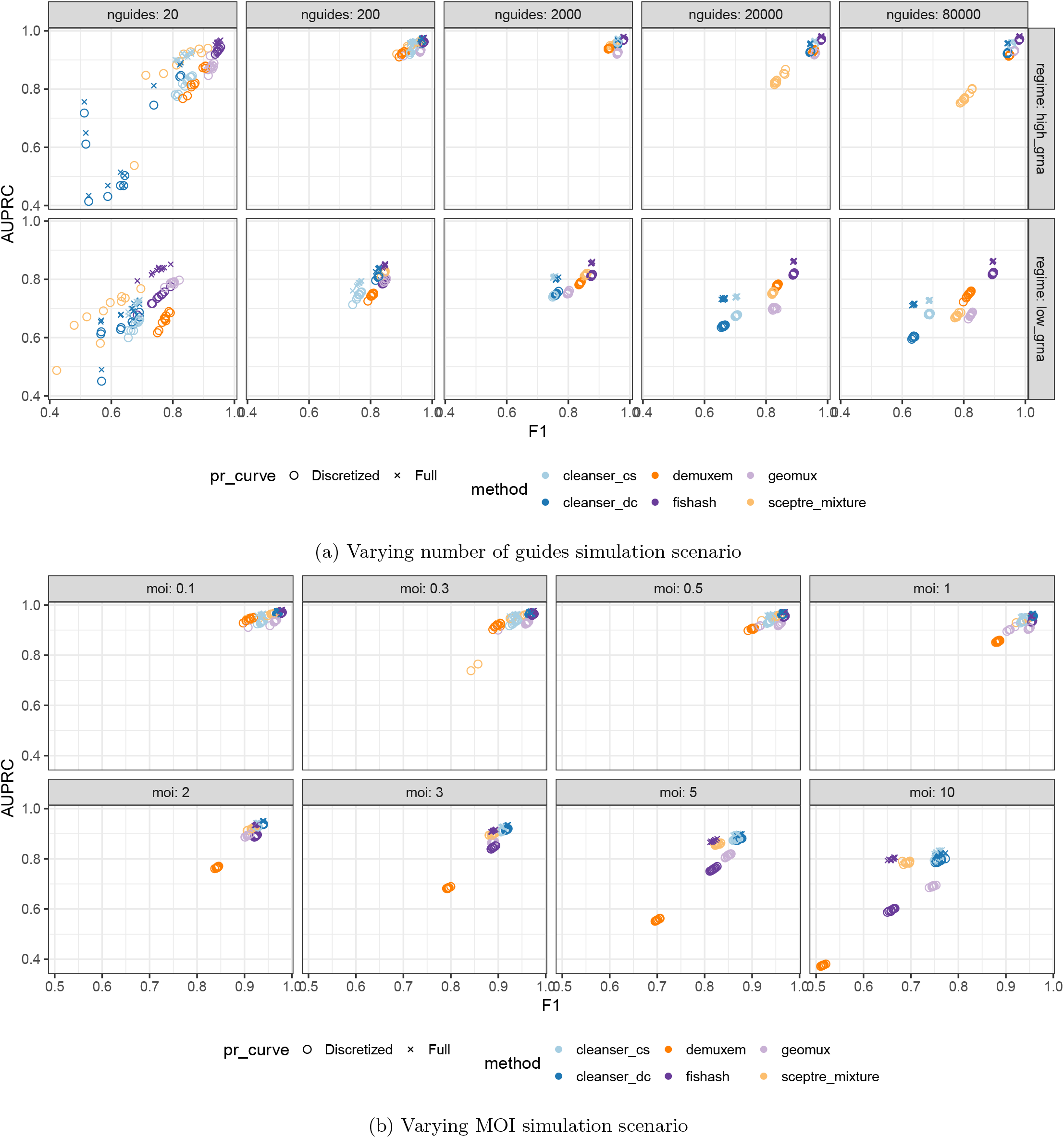
F1 scores at the default thresholds vs AUPRCs, in the two main simulation scenarios. In general, there is a qualitative concordance between the F1 scores and AUPRCs, though the interpretation is complicated due to the varying gap between the approximate (discretized) and exact (full) AUPRCs, the latter being only available for CLEANSER and fishash.

**Figure D.3.**
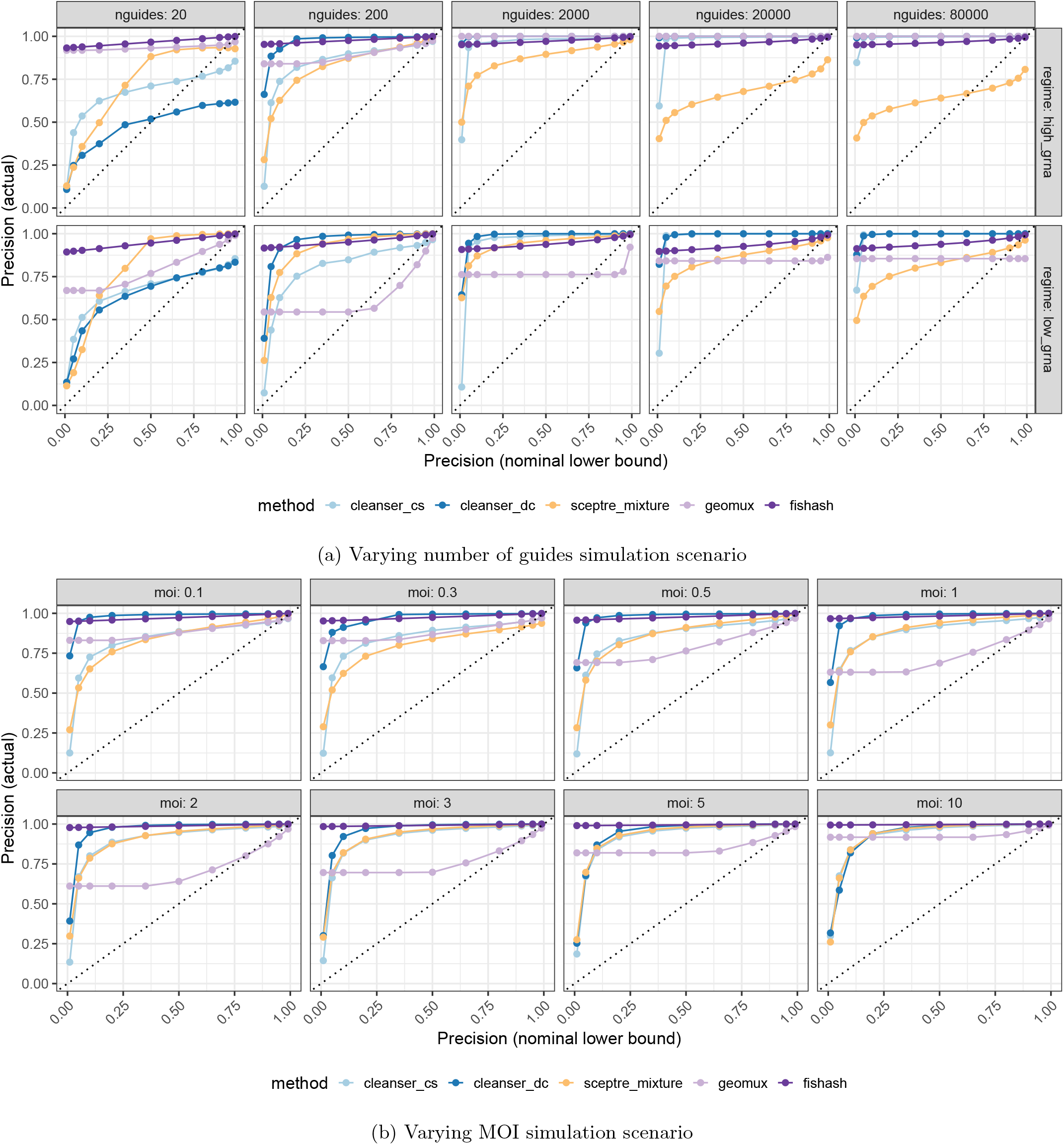
Precision and its nominal lower bound in discrete parameter sweep for main simulation scenarios. The precision nominal lower bound is the posterior probability cutoff for Bayesian methods (SCEPTRE, CLEANSER), or 1 minus the FDR cutoff for frequentist methods (fishash, geomux). The true precision was computed by aggregating the predictions from all 10 replicates per scenario. Fishash is conservative, and its precision remains high even as its nominal precision goes to 0. CLEANSER, SCEPTRE, and geomux all fail to control the FDR at least once (the precision drops below its nominal lower bound). DemuxEM was excluded because its tuning parameter does not correspond to a nominal precision.

**Figure D.4.**
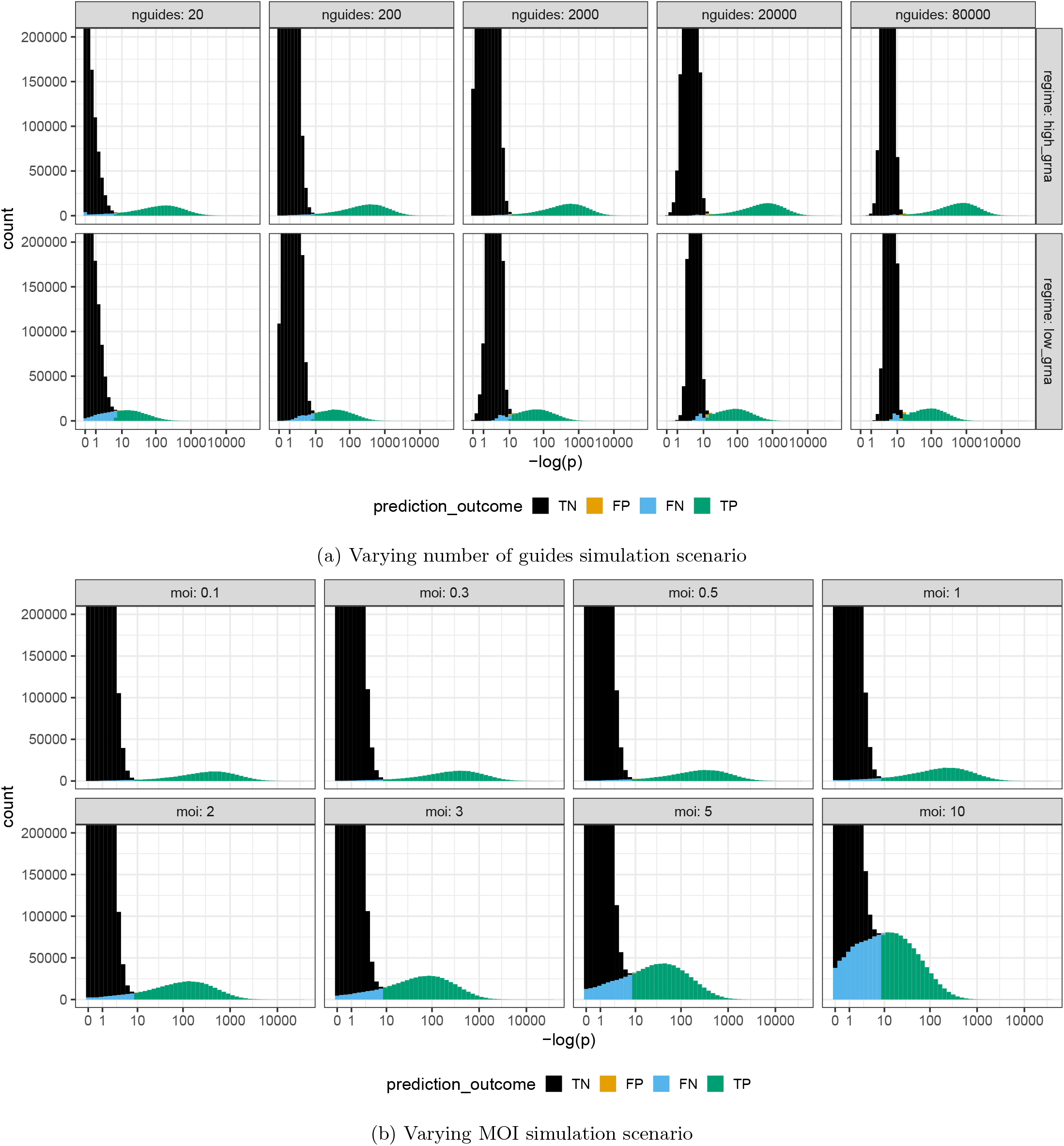
Fishash test statistic in main simulation scenarios, aggregated over replicates, colored by the prediction outcomes at the default FDR (0.05). One advantage of the fishash software is that it outputs this test statistic for all entries of the cell-guide matrix, allowing for visual selection of alternative cutoffs using these histograms, and for computing full precision-recall curves when benchmarking (Figures 5c, 6c).

## References

Barry, T., Roeder, K., and Katsevich, E. (2024). Exponential family measurement error models for single-cell crispr screens. Biostatistics, 25(4):1254–1272.

Barry, T., Wang, X., Morris, J. A., Roeder, K., and Katsevich, E. (2021). Sceptre improves calibration and sensitivity in single-cell crispr screen analysis. Genome biology, 22(1):344.

Benjamini, Y. and Hochberg, Y. (1995). Controlling the false discovery rate: a practical and powerful approach to multiple testing. Journal of the Royal statistical society: series B (Methodological), 57(1):289–300.

Benjamini, Y. and Yekutieli, D. (2001). The control of the false discovery rate in multiple testing under dependency. Annals of statistics, pages 1165–1188.

Braunger, J. M. and Velten, B. (2024). Guide assignment in single-cell crispr screens using crispat. Bioinformatics, 40(9):btae535.

Datlinger, P., Rendeiro, A. F., Schmidl, C., Krausgruber, T., Traxler, P., Klughammer, J., Schuster, L. C., Kuchler, A., Alpar, D., and Bock, C. (2017). Pooled crispr screening with single-cell transcriptome readout. Nature methods, 14(3):297–301.

Di Tommaso, P., Chatzou, M., Floden, E. W., Barja, P. P., Palumbo, E., and Notredame, C. (2017). Nextflow enables reproducible computational workflows. Nature biotechnology, 35(4):316–319.

Dixit, A. (2016). Correcting chimeric crosstalk in single cell rna-seq experiments. BioRxiv, page 093237.

Dixit, A., Parnas, O., Li, B., Chen, J., Fulco, C. P., Jerby-Arnon, L., Marjanovic, N. D., Dionne, D., Burks, T., Raychowdhury, R., et al. (2016). Perturb-seq: dissecting molecular circuits with scalable single-cell rna profiling of pooled genetic screens. cell, 167(7):1853–1866.

Fisher, R. A. (1922). On the interpretation of χ 2 from contingency tables, and the calculation of p. Journal of the royal statistical society, 85(1):87–94.

Fleming, S. J., Chaffin, M. D., Arduini, A., Akkad, A.-D., Banks, E., Marioni, J. C., Philippakis, A. A., Ellinor, P. T., and Babadi, M. (2023). Unsupervised removal of systematic background noise from droplet-based single-cell experiments using cellbender. Nature methods, 20(9):1323–1335.

Gaublomme, J. T., Li, B., McCabe, C., Knecht, A., Yang, Y., Drokhlyansky, E., Van Wittenberghe, N., Waldman, J., Dionne, D., Nguyen, L., et al. (2019). Nuclei multiplexing with barcoded antibodies for single-nucleus genomics. Nature communications, 10(1):2907.

Good, I. (1956). On the estimation of small frequencies in contingency tables. Journal of the Royal Statistical Society: Series B (Methodological), 18(1):113–124.

Guo, W. and Sarkar, S. (2020). Adaptive controls of fwer and fdr under block dependence. Journal of Statistical Planning and Inference, 208:13–24.

Hastie, T., Mazumder, R., Lee, J. D., and Zadeh, R. (2015). Matrix completion and low-rank svd via fast alternating least squares. The Journal of Machine Learning Research, 16(1):3367–3402.

Joag-Dev, K. and Proschan, F. (1983). Negative association of random variables with applications. The Annals of Statistics, pages 286–295.

Lehmann, E. (1966). Some concepts of dependence. The Annals of Mathematical Statistics, pages 1137–1153.

Liu, S., Hamilton, M. C., Cowart, T., Barrera, A., Bounds, L. R., Nelson, A. C., Dornbaum, S. F., Riley, J. W., Doty, R. W., Allen, A. S., et al. (2025). Characterization and bioinformatic filtering of ambient grnas in single-cell crispr screens using cleanser. Cell Genomics, 5(2).

Lopez, R., Regier, J., Cole, M. B., Jordan, M. I., and Yosef, N. (2018). Deep generative modeling for single-cell transcriptomics. Nature methods, 15(12):1053–1058.

Replogle, J. M., Saunders, R. A., Pogson, A. N., Hussmann, J. A., Lenail, A., Guna, A., Mascibroda, L., Wagner, E. J., Adelman, K., Lithwick-Yanai, G., et al. (2022). Mapping information-rich genotype-phenotype landscapes with genome-scale perturb-seq. Cell, 185(14):2559–2575.

Sarkar, T. K. (1969). Some lower bounds of reliability.

Sheng, C., Lopes, R., Li, G., Schuierer, S., Waldt, A., Cuttat, R., Dimitrieva, S., Kauffmann, A., Durand, E., Galli, G. G., et al. (2022). Probabilistic machine learning ensures accurate ambient denoising in droplet-based single-cell omics. BioRxiv, pages 2022–01.

Simpson, E. H. (1951). The interpretation of interaction in contingency tables. Journal of the Royal Statistical Society: Series B (Methodological), 13(2):238–241.

Teyssier, N. (2024). Computational Methods in Functional Genomics and Transcriptional Dynamics: Systems-Level Insights into Neurodegeneration and Neurodevelopmental Disorders. PhD thesis, University of California, San Francisco.

Yang, S., Corbett, S. E., Koga, Y., Wang, Z., Johnson, W. E., Yajima, M., and Campbell, J. D. (2020). Decontamination of ambient rna in single-cell rna-seq with decontx. Genome biology, 21(1):57.

Young, M. D. and Behjati, S. (2020). Soupx removes ambient rna contamination from droplet-based single-cell rna sequencing data. Gigascience, 9(12):giaa151.

Zappia, L., Phipson, B., and Oshlack, A. (2017). Splatter: simulation of single-cell rna sequencing data. Genome biology, 18(1):174.

